# Distinct chikungunya virus polymerase palm subdomains contribute to virus replication and virion assembly

**DOI:** 10.1101/2024.01.15.575630

**Authors:** Marie-France Martin, Boris Bonaventure, Nia E. McCray, Olve B. Peersen, Kathryn Rozen-Gagnon, Kenneth A. Stapleford

**Author notes:** Corresponding author: Kenneth A. Stapleford.

## Abstract

Alphaviruses encode an error-prone RNA-dependent RNA polymerase (RdRp), nsP4, required for genome synthesis, yet how the RdRp functions in the complete alphavirus life cycle is not well-defined. Previous work using chikungunya virus (CHIKV) has established the importance of the nsP4 residue cysteine 483 in maintaining viral genetic fidelity. Given the location of residue C483 in the nsP4 palm domain, we hypothesized that other residues within this domain and surrounding subdomains would also contribute to polymerase function. To test this hypothesis, we designed a panel of nsP4 variants via homology modeling based on the Coxsackievirus B3 3 polymerase. We rescued each variant in both mammalian and mosquito cells and discovered that the palm domain and ring finger subdomain contribute to polymerase host-specific replication and genetic stability. Surprisingly, in mosquito cells, these variants in the ring finger and palm domain were replication competent and produced viral structural proteins, but they were unable to produce infectious progeny, indicating a yet uncharacterized role for the polymerase in viral assembly. Finally, we have identified additional residues in the nsP4 palm domain that influence the genetic diversity of the viral progeny, potentially via an alteration in NTP binding and/or discrimination by the polymerase. Taken together, these studies highlight that distinct nsP4 subdomains regulate multiple processes of the alphavirus life cycle, placing nsP4 in a central role during the switch from RNA synthesis to packaging and assembly.

**Author Summary:** Chikungunya virus (CHIKV) is a re-emerging alphavirus transmitted to humans by mosquitoes and causing frequent explosive outbreaks. Its replication relies on a polymerase that incorporates a significant number of errors in the new genomes, making it a good candidate to develop vaccines or antiviral strategies. However, little is known on alphavirus polymerase function in alternate hosts. To begin to understand how the CHIKV polymerase nsP4 functions, we designed a panel of nsP4 variants taking advantage of the conservation of polymerase structure across positive strand RNA viruses. We discovered that the palm domain and ring finger of the polymerase were involved in host-specific RNA replication, genetic stability, and virus assembly. In addition, we demonstrated that the palm domain directly impacted the generation of viral genetic diversity. Taken together, these findings add further evidence to the crucial impact of the core palm domain of CHIKV polymerase not only on the replication of the RNA itself, but also on the genetic stability of the protein, as well as its involvement in viral assembly.

## Introduction

Chikungunya virus (CHIKV) is a re-emerging mosquito-borne virus, belonging to the *Togaviridae* family and the *Alphavirus* genus. Alphaviruses, which include both arthritogenic viruses like CHIKV and Mayaro virus (MAYV), and encephalitic viruses like Sindbis virus (SINV) and Venezuelan Equine Encephalitis virus (VEEV), can cause devastating disease and explosive outbreaks. Despite the severity of these diseases, there are limited vaccines or therapeutic strategies targeting alphaviruses. This fact, coupled with global climate change driving mosquito spread and the significant economic and social burden of alphaviral disease, highlights the need to better study how alphaviruses replicate at the molecular level (1–4).

CHIKV is an enveloped virus that possesses a ∼12 kb positive-sense single-stranded RNA genome flanked by a capped 5’-untranslated region (UTR) and a polyadenylated 3’-UTR. The genome contains two open-reading frames (ORF). The first ORF encodes for the nonstructural polyprotein P1234, that gives rise upon cleavage to the nonstructural proteins nsP1, nsP2, nsP3, and nsP4. The second ORF is under the control of an internal subgenomic promoter and encodes for the structural proteins capsid, E3, E2, 6K/TF and E1 (5). nsP1 is the methyl/guanylyltransferase responsible for capping new viral genomes (6), membrane curvature during replication complex formation (7) and anchoring of the replication complexes, also called spherules, at the plasma membrane (8,9). nsP2 has protease, helicase and NTPase activities (10–12) and controls fidelity in synergy with nsP4 (13). nsP3 possesses both a N-terminal ADP-ribose binding and hydrolase activities (14), as well as a hyper-variable C-terminal domain involved in binding cellular co-factors (15). Finally, nsP4 is the RNA-dependent RNA polymerase (RdRp) and possesses acetyltransferase activity (16), which is required to generate the polyadenylated tail at the 3’ end of the new viral genomes.

In addition to RNA polymerization, CHIKV nsP4 is a key contributor to the intrinsic viral fidelity as the cysteine 483 of nsP4 has been previously involved in the polymerase fidelity (13, 17,18). Fidelity is characterized by the ability of a polymerase to incorporate, based on a template, the correct nucleotide over a non-correct nucleotide during replication. Mutations at the position 483 of nsP4 result in varying phenotypes depending on the nature of the introduced residue. Whereas C483Y confers a high-fidelity phenotype to the polymerase (17), substitution with a glycine, C483G, results in a low-fidelity polymerase variant (18). Interestingly, both high- and low-fidelity polymerase variants are attenuated in mosquitoes and mice (17,18), highlighting the fine tuning of polymerase fidelity.

Recent work has unraveled the structures of alphavirus nsP4 and the replication complex (19,20). The elegant study by Tan *et al,* demonstrated that CHIKV active replication complex is composed of a dodecameric ring of nsP1 sitting at the neck of each spherule, at the center of which, one nsP2 molecule takes place on the cytoplasmic side, itself faced by one nsP4 molecule on the spherule side (19). Whereas these studies provide important structural insight on the CHIKV polymerase, nsP4 function is still not well-characterized. We suspect that the involvement of additional residues in nsP4 polymerization and fidelity remain to be elucidated.

In this study, we took advantage of the structural conservation of positive-sense RNA virus RdRp and the power of homology-based modeling to further understand CHIKV nsP4 function. Using the Coxsackievirus B3 (CVB3) 3 polymerase, a well-studied model for positive-sense RNA virus polymerase studies (4,5), we hypothesized that nsP4 residues in close vicinity to the active site of the polymerase, the GDD motif, and the previously characterized C483 residue could play important roles for polymerase function. To this end, we designed a set of 18 nsP4 variants targeting conserved residues among alphaviruses and investigated how these variants contribute to replication and virion production in mammalian and mosquito cells.

Using the full-length virus, we demonstrated that the CHIKV nsP4 palm domain and ring finger participate in host-specific replication and polymerase genetic stability. In addition, we demonstrated that introducing mutations in nsP4 affects viral assembly and impairs the generation of infectious progeny. Finally, we used deep-sequencing as an unbiased and high-throughput approach to elucidate how nsP4 variants impact the viral genetic diversity upon replication. We showed that mutating residues L383 and W486 in the palm domain and more specifically in close proximity to the C483 residue had important effect on the genetic diversity, presumably by affecting the binding and/or discrimination of NTPs during replication. Taken together, this study provides novel insight into nsP4 function, describing for the first time a role in viral assembly, and pinpoints important replication regulatory residues located in the palm domain of the polymerase.

## Results

### The CHIKV nsP4 palm and ring finger contribute to host-specific replication and polymerase genetic stability

Previous studies showed that the nsP4 residue C483, located in the palm domain (**Fig 1A**), played a key role for both polymerase fidelity and host-specific replication and virulence (13,17,18). We hypothesized that other nsP4 residues would also play important roles for nsP4 function and CHIKV life cycle. To test this hypothesis, we used homology modeling to design polymerase variants based on the Coxsackievirus B3 3D polymerase (CVB3 3D^pol^). As the crystal structure of the CHIKV polymerase has not been solved yet, we modeled nsP4 variants on the closely related alphavirus O’nyong’nyong viral protein which is 91% identical at the amino acid level (21) (**Fig. 1, S1 Fig**). This panel of CHIKV variants targets nsP4 residues conserved among alphaviruses (**Fig 1B**) including I312, L368 and T440 which correspond to known residues in CVB3 3D pol affecting polymerase fidelity (22,23), respectively I176, I230 and S299. In addition, we speculate that surrounding CHIKV nsP4 residues I372 (Y234 in CVB3 3D pol), L383 (W245 in CVB3 3D pol), L442 (I301 in CVB3 3D pol) and W486 (Y351 in CVB3 3D pol) could be involved in polymerase activity due to their respective orientations and locations, either in close vicinity to the GDD motif or the C483 residue (**Fig 1A**). A total of 18 nsP4 variants were designed by mutating 7 amino acids for further investigations.

**Fig 1.**
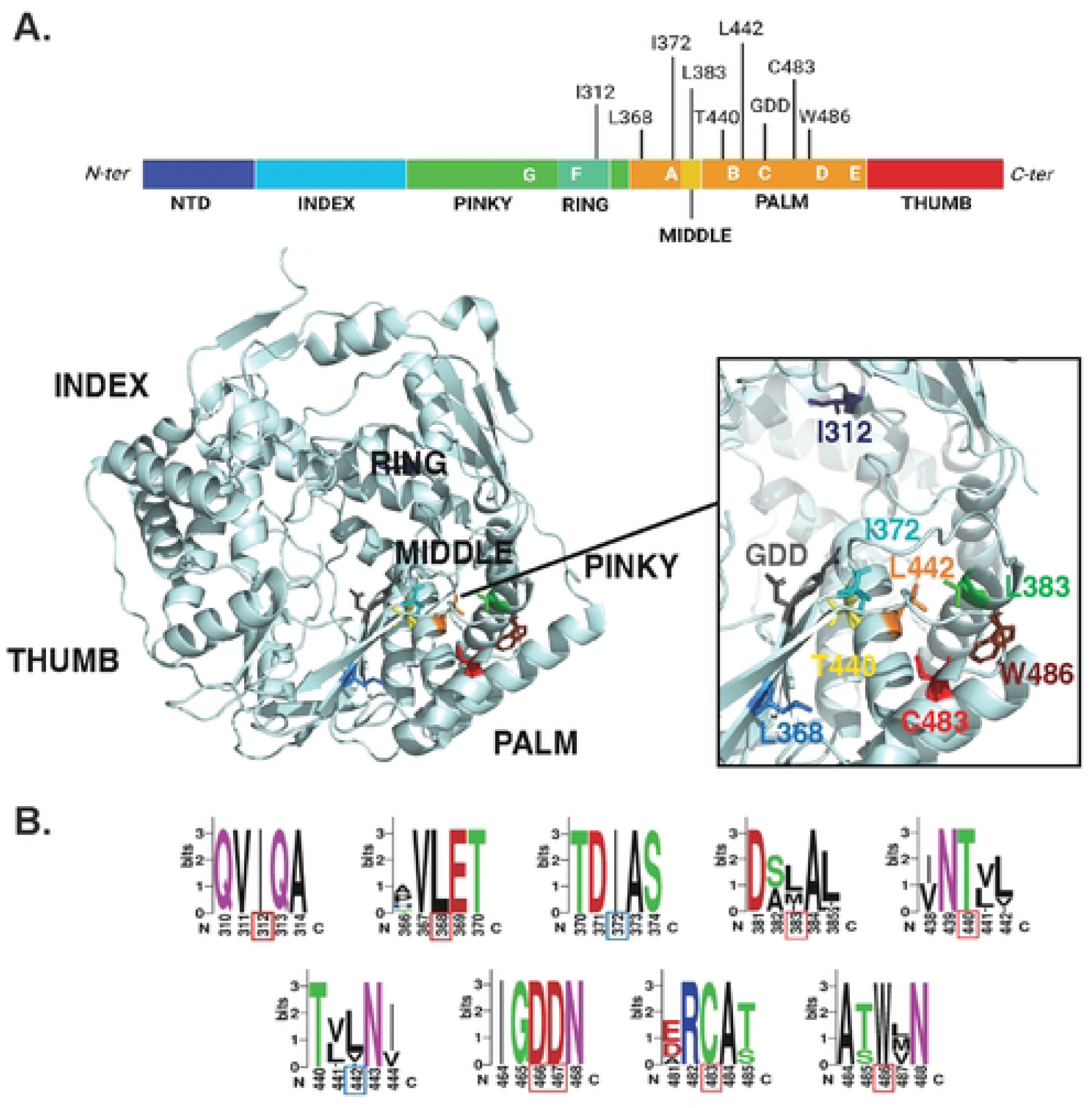
CHIKV nsP4 variant location and conservation. (**A**) Top: CHIKV nsP4 linear structure schematic colored from its N-terminus (dark blue) to its C-terminus (red). Subdomains of the protein are indicated below in capital letters and conserved positive-sense RNA virus polymerase motifs are delimited and indicated in white capital letters onto each subdomain. Each mutated residue is indicated at the top of the protein with a tick. Schematic made with BioRender. Bottom: O’nyong’nyong (ONNV) nsP4 3D structure (PDB: 7Y38) annotated with subdomains (black). Mutated residues displayed on nsP4 3D structure with their side chain color-coded as follow from their N-ter to C-ter position in nsP4: I312 (dark blue), L368 (medium blue), I372 (turquoise), L383 (medium green), T440 (yellow), L442 (orange), GDD motif (grey), C483 (red), W486 (brown). (**B**) WebLogo of mutated residues among alphaviruses generated using the multiple sequence alignment of nsP4 via MUSCLE (EMBL-EMI; see Methods). The height of each stack of letters represents the sequence conservation at the indicated position and the height of each symbol within the stack represents its relative frequency. A patch of five amino acids total was selected, with each mutated residue at the center of it framed by a red square (if outside of a conserved motif) or blue (if inside a conserved motif). Each color represents amino acids of similar properties.

As CHIKV replicates both in mammals and mosquitoes in nature, we wanted to investigate the impact of each nsP4 variant on CHIKV replication in both hosts (**Fig 2**). To do so, we selected as model cell lines, the hamster BHK-21 cells for the mammalian host, and the *Aedes albopictus* C6/36 clone cell line for the mosquito host. First, we tried to rescue each nsP4 variant by transfecting the *in vitro* transcribed full-length CHIKV nsP4 variant RNA into BHK-21 or C6/36 cells. This approach has the advantage to synchronize replication and eliminate any confounding effects of virus entry. At 48 hours post transfection (hpt), we titered the supernatant by plaque assay on Vero cells to determine infectious viral particles production, quantified extracellular CHIKV genomic RNA by RT-qPCR, and Sanger sequenced the nsP4 variant to address genetic stability.

**Fig 2.**
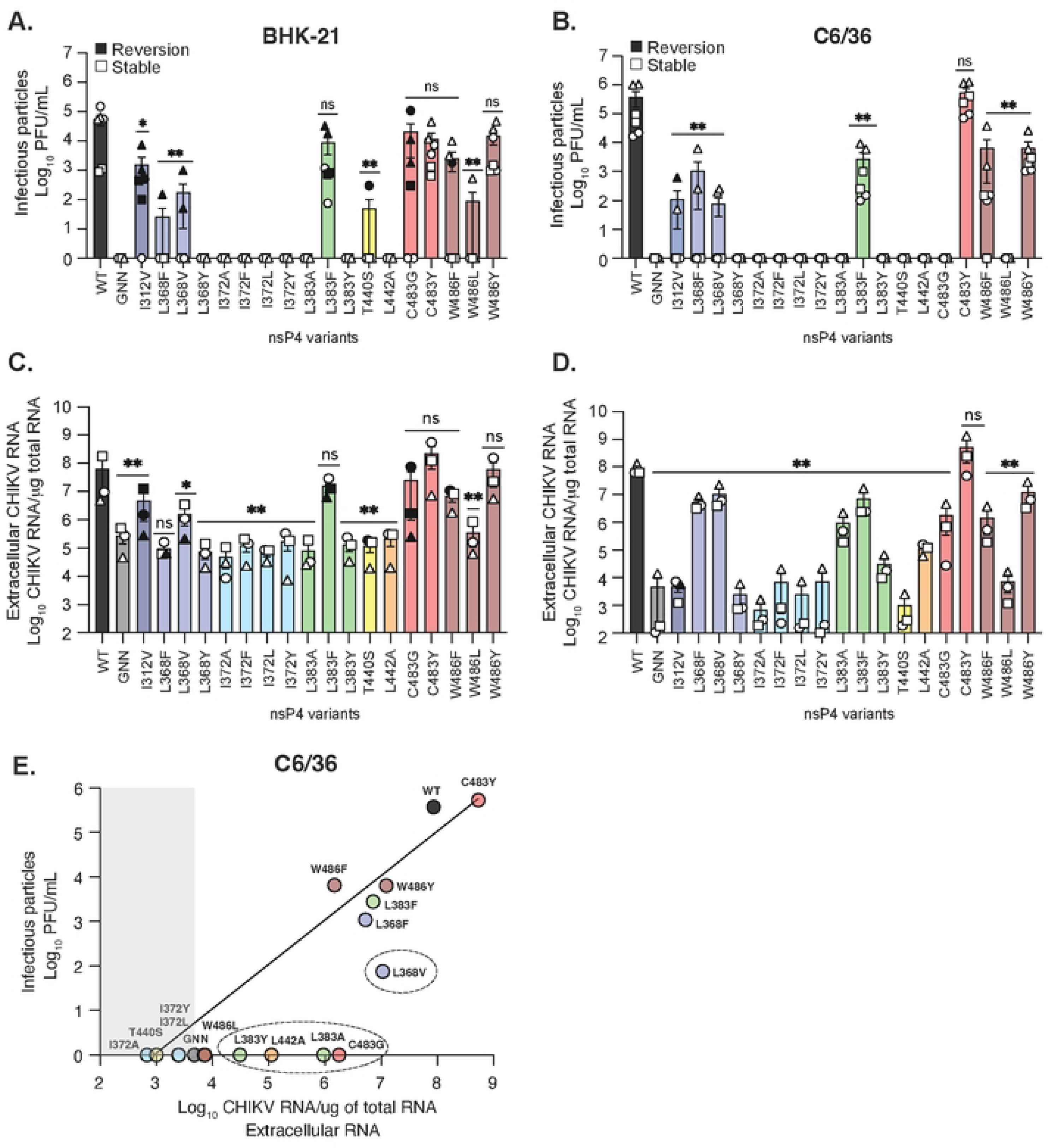
CHIKV nsP4 variants replicate in a host-specific fashion. BHK-21 (left side) or C6/36 (right side) cells were transfected in a 6-well plate with *in vitro* transcribed full-length CHIKV RNA of each variant and incubated for 48 hours at 37°C and 28°C respectively. (**A and B**) Culture supernatants of each condition were harvested at 48 hpt and infectious virus was quantified by plaque assay on Vero cells. (**C and D**) RNA was extracted from culture supernatants and CHIKV genomes were quantified by RT-qPCR. Genetically stable or reverted nsP4 variants are respectively depicted with a solid black symbol or an empty symbol on **A to D**. (**E**) A linear regression analysis was performed on infectious viral particles and extracellular viral genomic RNA for C6/36 cells. A grey shade is highlighting the nsP4 variants behaving like the negative control, nsP4 GNN, and dashed circles are pointing to clusters of similar variants. For all panels, each symbol represents an independent biological experiment, in technical transfection duplicate, with solid symbols showing the nsP4 variants that reverted to the WT residue and clear symbols the variants that did not revert. A Mann-Whitney *t* test was performed against nsP4 WT for all data and data with statistical significance was labeled as follow: * P<0.05, ** P<0.01. Graphs show the average and SEM of three independent experiments with internal transfection duplicates on all panels.

In BHK-21 cells (**Table 1**, **Fig 2A and C**), nine out of eighteen nsP4 variants produced infectious virus (I312V, L368F, L368V, L383F, T440S, C483G, C483Y, W486F and W486Y). Virus production (**Fig 2A**) was correlated with the presence of extracellular RNA (**Fig 2C**), while nsP4 variants that did not produce infectious virus showed viral RNA levels similar to the replicase-dead GNN variant, used as a negative control (**Fig 2C**). However, the sequencing of the nsP4 region in CHIKV genomes from the rescued variants showed that only three of them, C483Y, W486L and W486Y, were genetically stable (**Table 1**). Every other nsP4 rescued variant showed either a partial or a total reversion to the WT residue at 48 hpt (depicted as a solid symbol in **Fig 2**). It is likely that the variants that reverted had a decreased fitness, resulting in the selection of revertant virus over the course of the rescue. By looking at the specific codon changes during reversion, we noticed that the majority of revertant viruses presented only one nucleotide change compared to nsP4 WT with the exception of nsP4 W486F (**Table 1**). Indeed, this variant reverted by changing two nucleotides, from UUC (F) to UGG (W) with tryptophan being encoded by a unique UGG/TGG codon. Surprisingly, although nsP4 L368F and nsP4 L368V both reverted in BHK-21 cells, the reversion was achieved through two different codons, respectively CUU and UUG, suggesting potentially less stringent codon usage at residue 368.

**Table 1.**

| nsP4<br>variant | % of replicates<br>being rescued |  | Genetic stability |  | % of revertant<br>amongst rescued<br>viruses |  | Reversion genotype |  |
| --- | --- | --- | --- | --- | --- | --- | --- | --- |
|  | BHK-21 | C6/36 | BHK-21 | C6/36 | BHK-21 | C6/36 | BHK-21 | C6/36 |
| I312V | 83 | 33 | Reverted to I | Reverted to I | 100 | 50 | GTA (V) to<br>ATA (I) | GTA (V) to<br>ATA (I) |
| L368F | 16 | 33 | Reverted to L | Stable | 100 | 0 | TTT (F) to<br>CTT (L) | Stable |
| L368V | 33 | 33 | Reverted to L | Stable | 100 | 0 | GTG (V) to<br>TTG (L) | Stable |
| L368Y | 0 | 0 | Not rescued | Not rescued | N/A | N/A | N/A | N/A |
| I372A | 0 | 0 | Not rescued | Not rescued | N/A | N/A | N/A | N/A |
| I372F | 0 | 0 | Not rescued | Not rescued | N/A | N/A | N/A | N/A |
| I372L | 0 | 0 | Not rescued | Not rescued | N/A | N/A | N/A | N/A |
| I372Y | 0 | 0 | Not rescued | Not rescued | N/A | N/A | N/A | N/A |
| L383A | 0 | 0 | Not rescued | Not rescued | N/A | N/A | N/A | N/A |
| L383F | 100 | 100 | Reverted to L | Stable | 66 | 0 | TTT (F) to<br>CTT (L) | Stable |
| L383Y | 0 | 0 | Not rescued | Not rescued | N/A | N/A | N/A | N/A |
| T440S | 16 | 0 | Reverted to T | Not rescued | 16 | N/A | TCA (S) to<br>ACA (T) | N/A |
| L442A | 0 | 0 | Not rescued | Not rescued | N/A | N/A | N/A | N/A |
| C483G | 66 | 0 | Reverted to C | Not rescued | 66 | N/A | GGT (G) to<br>TGT (C) | N/A |
| C483Y | 100 | 100 | Stable | Stable | 0 | 0 | Stable | Stable |
| W486F | 50 | 83 | Reverted to W | Stable | 33 | 0 | TTC (F) to<br>TGG (W) | Stable |
| W486L | 16 | 0 | Stable | Not rescued | 16 | N/A | Stable | N/A |
| W486Y | 100 | 100 | Stable | Stable | 0 | 0 | Stable | Stable |

N/A : non applicable

In mosquito cells (**Table 1**, **Fig 2B, D, and E**), seven variants produced infectious virus (I312V, L368F, L368V, L383F, C483Y, W486F and W486Y). As we observed in BHK-21 cells, extracellular CHIKV RNA correlated with the presence of infectious virus in C6/36 cells (**Fig 2D**). However, in these cells nsP4 variants L368V, L383A, L383Y, L442A, as well as C483G released extracellular viral RNA (**Fig 2D**) in the absence of infectious viral particles (**Fig 2B**), suggesting these variants are able to synthesize RNA, yet not form infectious particles. It must be noted that no CPEs was observed in these conditions at the moment of the harvest, contrary to the nsP4 WT, L383F, C483Y and W486Y conditions (data not shown), excluding the possibility of intracellular content release upon cell death. Importantly, sequencing revealed that six rescued variants were genetically stable with the exception of nsP4 I312V which reverted once upon two rescue events in this cell type (**Table 1**). To better visualize the results from C6/36 cells, we correlated infectious viral particle production and extracellular viral RNA release in mosquito cells (**Fig 2E**) and highlighted genetically stable variants with no infectious particles but high levels of extracellular RNA (**Fig 2E, dashed circles**).

Together, these results reinforce host species greatly influence CHIKV variant replication and genetic stability. Mosquito cells tend to provide a less constrained environment for polymerase activity, while a mammalian host seems to be applying a more stringent selection. Moreover, these studies highlight critical residues involved in polymerase function as no variant of the I372 residue produced infectious virus in either cell type (**Table 1**, **Fig 2A and B**), even if substitutions were made with other comparable hydrophobic amino acids. This suggests the presence of an important selective pressure denoting an essential role for an isoleucine at position 372 in RdRp activity across species. Similarly, neither of the nsP4 variants L368Y, L383A, L383Y, or L442A produced infectious virus in either cell type (**Table 1**, **Fig 2A and B**). However, contrary to nsP4 L383A and L383Y, the L383F variant did produce viral progeny despite the close similarity between these two amino acid substitutions and the conservation of a hydrophobic residue at this position (**Fig 1B**).

### The nsP4 palm domain contributes to infectious particle production

Given that in C6/36 cells several of the CHIKV nsP4 variants led to no infectious particles production but release high levels of extracellular RNA (**Fig 2**), we wanted to confirm that these nsP4 variants were capable to replicate in the cells by assessing intracellular viral RNA and protein levels (**Fig 3**). Viral RNA was quantified by RT-qPCR on total RNA extracts from transfected cells corresponding to Figure 2. In mammalian cells, only the C483Y and W486Y nsP4 variants, which were also the ones that did not revert to the WT residue, had similar levels of intracellular RNA to WT CHIKV (**Fig 3A**). The other nsP4 variants showed similar viral RNA levels to the negative control, nsP4 GNN, with nsP4 L383F and C483G showing an in-between phenotype, but reverting to WT CHIKV. Intracellular viral protein levels were assessed by immunoblot for nsP4 and the structural proteins capsid and E2 at 48 hpt from the supernatants of Figure 2. Overall, in BHK-21 cells (**Fig 3C, S2A and B**), the presence of nsP4, capsid and E2 correlated with intracellular RNA levels, yet again many of these variants reverted to the WT residues (**red stars on Fig 3C, S2A and B**). Interestingly, the nsP4 variant W486F, which did not revert in this experiment, led to WT levels of nsP4, less capsid production but had an even lower level of E2 accumulation. This could be explained by the chronologically differential processing of capsid and the envelope glycoproteins, capsid being processed before E2 during the replication cycle, thus suggesting that nsP4 W486F is likely delayed in its replication comparing to nsP4 WT.

**Fig 3.**
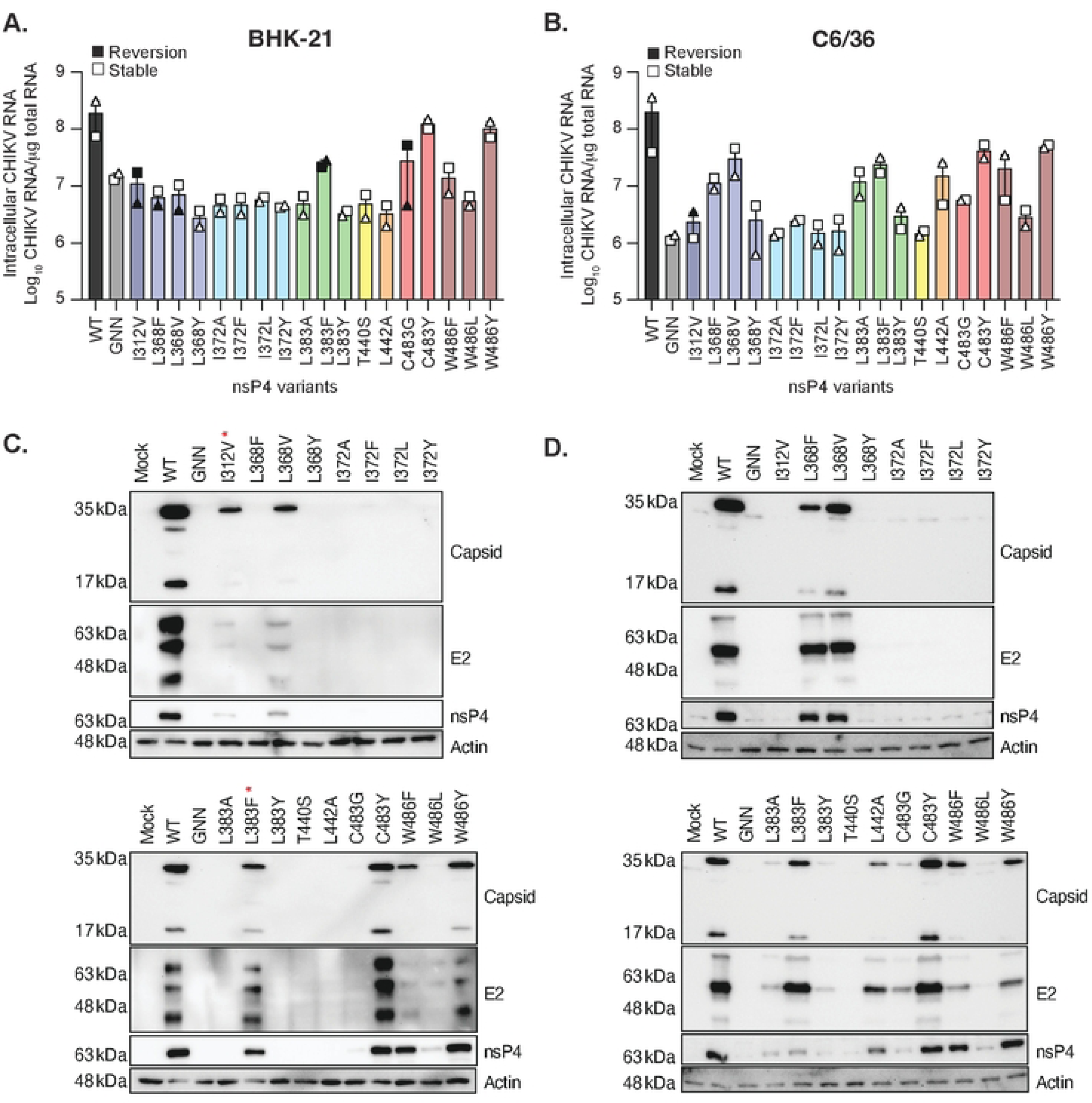
Intracellular RNA and protein accumulation of nsP4 variants in mammalian and mosquito cells. The cells from each technical duplicate transfections in Figure 2 were used for these analyses. BHK-21 cells (left side) or C6/36 cells (right side) cells were transfected in a 6- well plate with *in vitro* transcribed full-length CHIKV nsP4 variants and incubated for 48 hours at 37°C and 28°C respectively. (**A and B**) Cells were harvested, and intracellular RNA purified to quantify the amount of CHIKV RNA per μg of total RNA by RT-qPCR. Each symbol represents an independent biological experiment with solid symbols showing the nsP4 variants that reverted to the WT residue and clear symbols the variants that didn’t revert. A Mann-Whitney *t* test was performed against nsP4 WT and data with statistical significance was labeled as follow: *P<0.05, ** P<0.01. Graphs show the average and SEM of two independent experiments in technical duplicates. (**C and D**) Cells were transfected with WT CHIKV or each variant *in vitro* transcribed RNA as above (and corresponding to Fig 2) and were lysed at 48 hpt for SDS-PAGE and immunoblotting for the structural proteins Capsid and E2 (top of a membrane), nsP4 (middle of a membrane) and the house-keeping gene actin (bottom of a membrane) with molecular weights on the left side as a reference. The nsP4 variants were split onto two distinct membranes, ran in the same tank, with all three controls, Mock, nsP4 WT and nsP4 GNN, were ran on each gel and the nsP4 variants organized from the N-ter (left of the membrane) to the C- ter (right of the membrane) of nsP4. Membranes are representative of three independent experiments. The red star highlights the nsP4 variants that reverted in the specific experiment shown here. The western blots of other experiments can be found in Fig S2 and S3.

In mosquito cells, most of the genetically stable CHIKV nsP4 variants we have been able to rescue had intracellular viral RNA levels above the nsP4 GNN control (**Fig 3B**), including the nsP4 variants, L368F, L368V, L383A, and L442A. Of these variants, nsP4 L383A and L442A were associated with an absence of infectious viral particle release, and the nsP4 variants L368F and L368V with a low level of infectious progeny when rescued (**Fig 2B**). This indicates that these variants are replication competent but defective in virion production. As in mammalian cells, the presence of intracellular viral proteins was assessed in mosquito cells with an immunoblot against nsP4, capsid and E2. Importantly, we observed the production of nsP4, capsid, and E2 by nsP4 variants that did not produce consistently infectious particles, namely nsP4 L368F, L368V, L383A, L383Y, L442A and C483G (**Fig 3D, S3A and B**). Taken together these results show that while nsP4 variants are not making infectious particles, they are producing viral nonstructural and structural proteins, suggesting an impairment in viral production. This indicates a role for nsP4 in the proper assembly or secretion of infectious particles and emphasize again the impact of the host specie on viral replication.

### CHIKV nsP4 variants L368F, L368V and L442A have impaired infectious progeny release and intracellular protein accumulation

Failing of producing viral particles may be explained by a delay in virion production. To assess if the nsP4 variants that do not produce viral particles had delayed replication, we performed growth curves in C6/36 cells (**Fig 4**), focusing on the nsP4 variants that produce few or no infectious particles (L368F, L368V, L383A, L383F, L383Y, L442A, and C483G). Cells were transfected with the nsP4 variant *in vitro* transcribed full-length RNA, culture supernatants were harvested at 24, 48, and 72 hpt, and infectious viral particles were titered by plaque assay. We observed that nsP4 variants L368F (**Fig 4A**), and L383F (**Fig 4B**) were delayed or attenuated in virion production with kinetics comparable to the nsP4 variant C483G (**Fig 4C**), which is known to be attenuated in insect cells (18). Interestingly, the CHIKV nsP4 variants L368V (**Fig 4A**), L383A (**Fig 4B**), L383Y (**Fig 4B**), and L442A (**Fig 4C**) did not produce infectious particles even at the late time point of 72 hpt, indicating the lack of virion production was not due to a replication delay.

**Fig 4.**
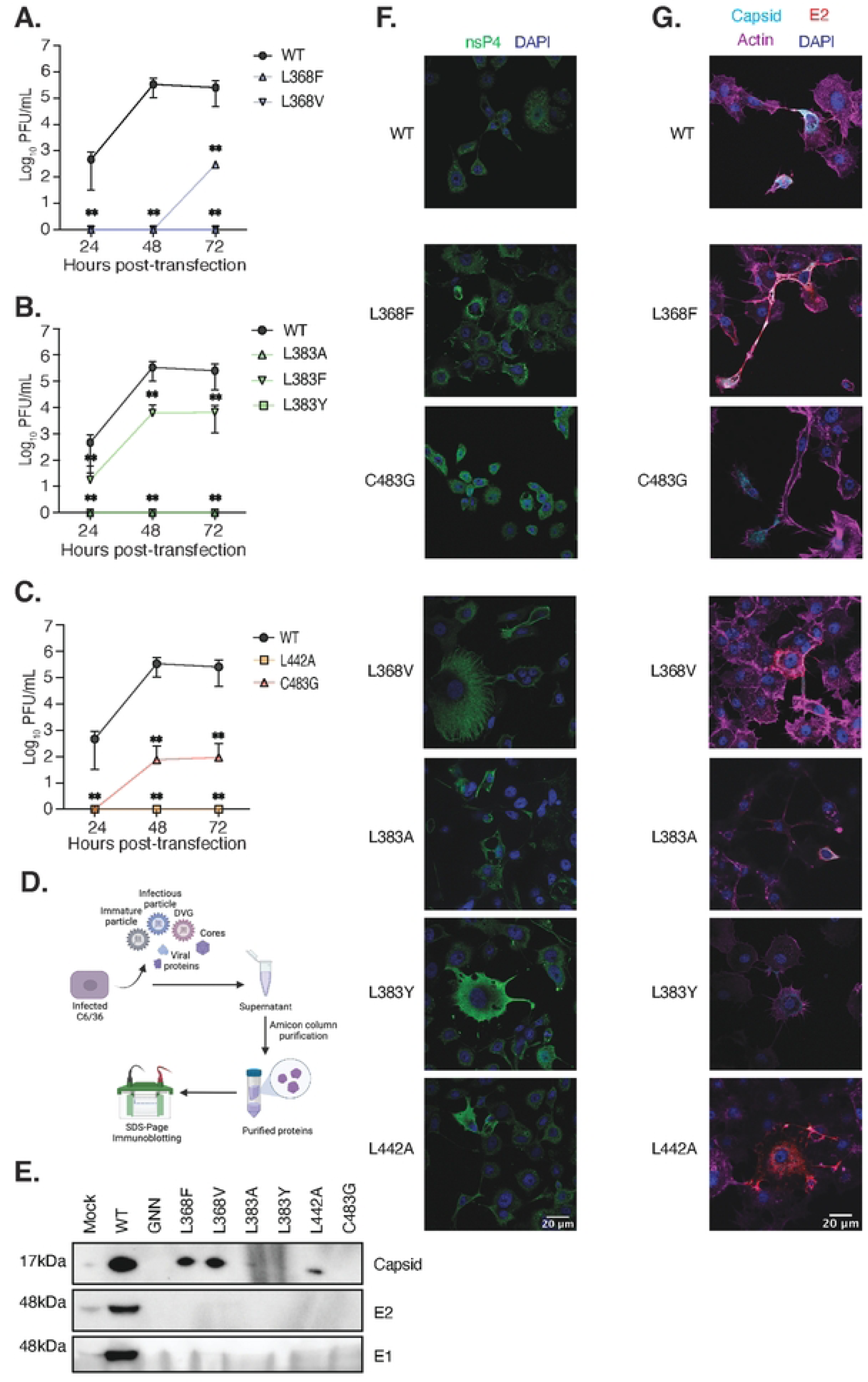
Growth curve and cellular viral protein expression of defective nsP4 variants in mosquito cells. C6/36 cells were transfected in a 24-wll plate with *in vitro* transcribed full-length CHIKV genetically stable nsP4 variants and incubated at 28°C. Supernatants were taken at 24hpt, 48 hpt and 72 hpt and infectious particles were quantified on Vero cells by plaque assay (**A, B, and C**). Each panel shows all the nsP4 variants corresponding to the same residue with nsP4 WT (black) as a positive control. For all panels, a Mann-Whitney *t* test was performed against nsP4 WT and data with statistical significance was labeled as follow: * P<0.05, ** P<0.01. Graphs show the average and SD of three independent experiments with internal transfection duplicates on all panels. (**D and E**) Supernatants from C6/36 cells transfected with *in vitro* transcribed full-length CHIKV nsP4 variants that did not show any infectious viral particles, but intracellular viral proteins were harvested at 48 hpt (corresponding to Fig 2). Extracellular proteins were purified with Amicon columns (**D**, schematic made with BioRender). Denatured proteins were subjected to a western blotting (**E**) and analyzed for the presence of the structural proteins capsid (top), E2 (middle) and E1 (bottom). (**F and G**) C6/36 cells were seeded onto coverslips and transfected with *in vitro* transcribed full-length CHIKV nsP4 variants then incubated for 24 hours at 28°C. Cells were fixed and stained nsP4 (green) and DAPI (blue) or for Capsid (light blue), E2 (red), Actin (purple) and DAPI (blue) respectively (**D**) and (**E**) and images acquired on a confocal microscope. nsP4 variants were sorted between the control (WT), the variants that have delayed replication (L368F and C483G) and the abortive variants (L368V, L383A, L383Y and L442A). Images are representative of two independent experiments for all panels. Scale bar: 20 Ιm. Images of negative control mock and nsP4 GNN transfected cells as well as additional nsP4 variant images can be found in Fig **S5**.

Although several variants do not produce infectious virus, it is possible they secrete noninfectious particles. To test the hypothesis of a noninfectious viral structure release we used the supernatants from mosquito cells transfected with the nsP4 variants L368F, L368V, L383A, L383Y, L442A and C483G at 48hpt in Figure 2, and concentrated them with Amicon Ultra centrifugal filters (**Fig 4F**). We then addressed the presence of structural proteins capsid, E1 and E2 by immunoblotting (**Fig 4G**, **S4A and B**). Interestingly, capsid protein was detected for the nsP4 variants L368F, L368V and L442A, alongside nsP4 WT (**Fig 4G**). Nevertheless, no E1 or E2 could be recovered in this specific experiment, whereas a small amount of E2 protein appears for nsP4 L368V in another independent experiment (**Fig S4A**). This suggest that nsP4 variants L368F, L368V and L442A produce noninfectious particles containing the viral capsid (**Fig 4G**) and likely viral RNA as we detected in **Fig 2D**. The detection of intracellular viral structural proteins as shown in **Fig 3D** and **Fig 4E** exclude the possibility that these nsP4 variants fail to express the viral glycoprotein. These observations are of special importance as they point towards a role of CHIKV polymerase in viral assembly, likely via interaction of the newly synthesized genomes with capsid

Finally, to explore any potential issues with nsP4 variant structural protein expression and subcellular localization in C6/36 cells, we performed immunofluorescence at 24 hpt on each assembly defective variant, staining for nsP4 (**Fig 4D**) and capsid, E2, and actin (**Fig 4E, Fig S5**). Staining of uninfected mock and nsP4 GNN negative controls are shown in **Fig S5**. For all the attenuated nsP4 variants (L368F, C483G, W486F and W486Y) or the ones which do not produce infectious particles (L368V, L383A, L383Y and L442A), intracellular polymerase expression was similar to nsP4 WT (**Fig 4D**). However, the nsP4 variants L368V and L383Y nsP4 presented as a diffuse staining in contrast to punctate nsP4 staining for the nsP4 WT virus. Looking then at structural proteins and actin expression (**Fig 4E, Fig S5C**), we noted several interesting observations. First, nsP4 WT, L368F, L368V, and L383A all expressed capsid and E2 proteins, even if with some difference in intensity correlating with the western blot (**Fig 3D**). Interestingly, nsP4 C483G expressed capsid, but no E2 glycoprotein. This is in line with the delayed replication observed in **Fig 4C**, which is in alignment with the molecular biology of CHIKV where E2 protein is processed after capsid during the viral replication cycle. On the other hand, at 24hpt the nsP4 variant L383Y do not express capsid or E2 (**Fig 4E, Fig S5C**), an observation that correlate with the absence of infectious particles being released even at a late time-point such as 72 hpt (**Fig 4B**). Finally, we observed an important expression of E2 but very few of capsid for the nsP4 variant L442A at 24hpt (**Fig 4E, Fig S5C**), whereas both proteins are correctly expressed at 48hpt (**Fig 3D, Fig S3**). Coupling these observations with actin fibers expression, we could gain insight into the potential biology of the nsP4 variants. Indeed, nsP4 L368F and L368V showed an actin expression pattern comparable to the non-infected cells, with a notable actin accumulation at the junction of infected cells with either infected or non-infected cells in the case of L368F, similarly to nsP4 WT (**Fig 4E, Fig S5C**). Of note, it seems that both nsP4 variants, as nsP4 WT, were able to produce intercellular extensions that have been described as actin and tubulin positive and of more than 10 µm long (24) (see scale bar on **Fig 4E, Fig S5**). These CHIKV-induced extensions have been recently shown to be required for cell-to-cell viral spread and to participate in CHIKV’s ability to escape circulating neutralizing antibodies (25). In addition, they require the presence of both capsid and E2 proteins for their formation. Interestingly, the F-actin network appears to be disorganized for the nsP4 variants L383A and L442A, and its expression less important for the nsP4 variant L383Y (**Fig 4E, Fig S5C**). A disorganization of the F-actin network could potentially be linked to a defect in the transport of E2 towards the plasma membrane or other viral assembly step.

Putting together these observations with their growth curves, we speculate that the poor actin network could be linked to the failure to produce infectious viral progeny, as actin is known to be important for viral assembly.

### Residues in the nsP4 C483 pocket affect virus growth, genetic diversity of the progeny, and the substitution profile of the polymerase

By having an intrinsic error rate, RdRps are generating viral diversity while maintaining genomic integrity. The CHIKV nsP4 residue C483 resides in the polymerase palm subdomain and has been shown to regulate virus fidelity and host specific growth (17,18). Given that several of the nsP4 residues we rescued and were genetically stable (L383 and W486) are in close proximity to residue C483, we hypothesized these residues would also be important for viral growth and genetic diversity.

First, we addressed virus growth kinetics as described previously (**Fig 5**). In BHK-21 cells (**Fig 5A and B**), the dead-replicase nsP4 GNN did not produce viral progeny, the nsP4 C483Y variant replicated similarly to nsP4 WT, and the nsP4 W486Y variant showed delayed replication, with infectious viral particles not being released until 48 hpt (**Fig 5A**). To address whether viral proteins were correctly expressed and localized during infection, we looked by immunofluorescence at CHIKV nsP4 cellular expression and localization at 24 hpt (**Fig 5B**). CHIKV nsP4 WT, C483Y, and W486Y displayed a punctuated cytoplasmic nsP4 expression suggesting that nsP4 was behaving similarly between the variants. These results indicate that the CHIKV nsP4 W486Y variant has delayed growth kinetics in BHK-21 cells, at least not due to an altered expression or subcellular localization of nsP4. In C6/36 cells (**Fig 5C and D**), we found that the C483Y variant grew like WT CHIKV, yet W486F and W486Y viral productions were delayed and produced viral progeny only at 48 or 72 hpt (**Fig 5C**). These results are in line with the delayed growth of the nsP4 L383F variant in C6/36 cells (**Fig 4B**). When we looked at nsP4 expression by immunofluorescence, we found that the C483Y variant behaved like WT CHIKV while the nsP4 W486Y variant showed more cytoplasmic staining (**Fig 5D**).

**Fig 5.**
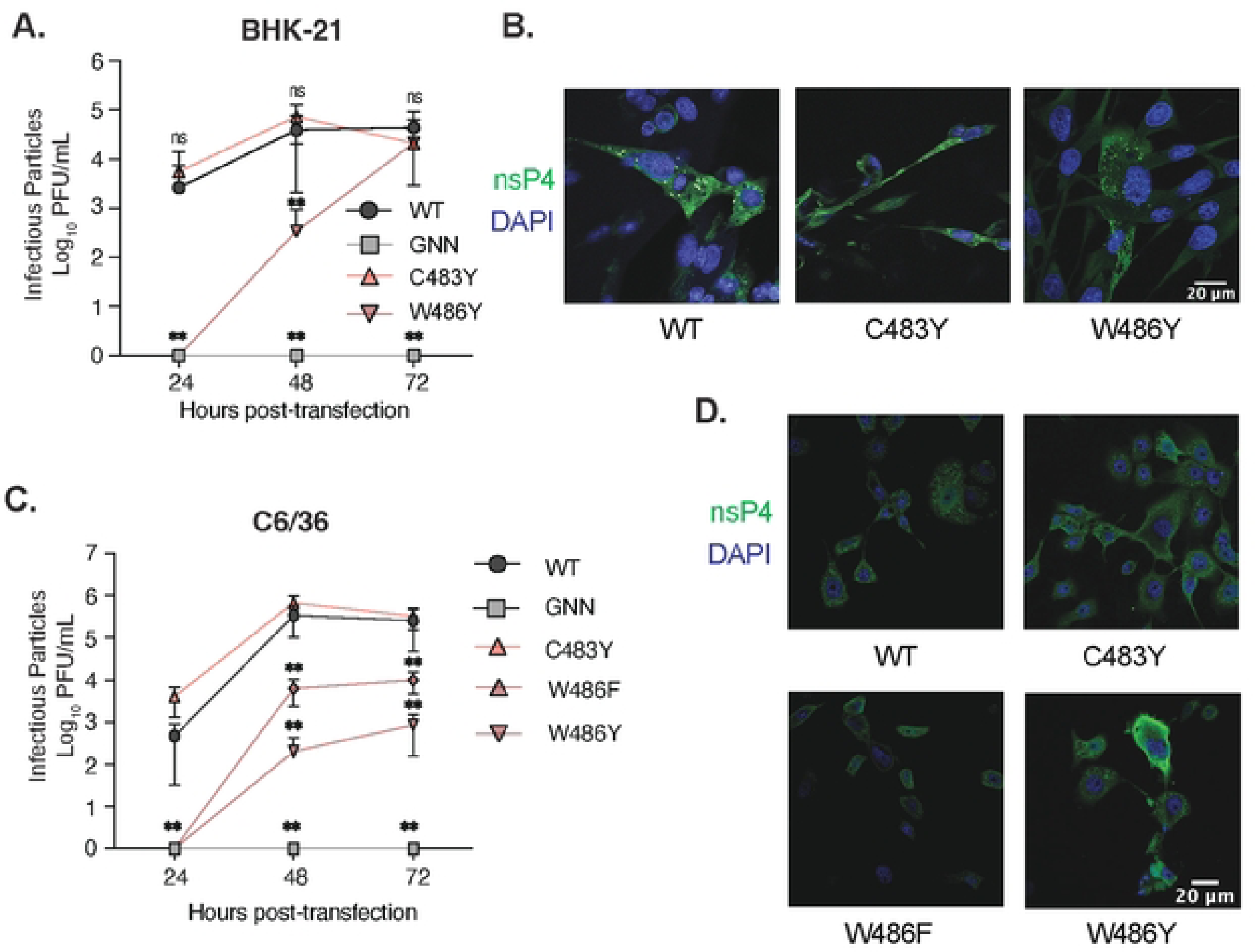
Growth curve and cellular nsP4 expression of stable nsP4 variants in mammalian cells and mosquito cells. BHK-21 cells (**A and B**) or C6/36 cells (**C and D**) were transfected in a 24- well plate with *in vitro* transcribed full-length CHIKV genetically stable nsP4 variants and incubated at 37°C and 28°C respectively. Supernatants were taken at 24hpt, 48 hpt and 72 hpt and infectious particles quantified on Vero cells by plaque assay (**A and C**). Each panel show the nsP4 variants corresponding with both nsP4 WT (black) and nsP4 GNN (grey) as controls. For all panels, a Mann-Whitney *t* test was performed against nsP4 WT an2d data with statistical significance was labeled as follow: * P<0.05, ** P<0.01. Graphs show the average and SD of three independent experiments with internal transfection duplicates on all panels. BHK-21 cells (**B**) or C6/36 cells (**D**) were seeded onto coverslips and transfected with *in vitro* transcribed full-length CHIKV nsP4 variants then incubated for 24 hours at 37°C. Cells were fixed and stained for nsP4 (green) and DAPI (blue) and images acquired on a confocal microscope. Images are representative of two independent experiments for all panels. Scale bar: 20 Ιm.

To assess the impact of polymerase variants on the genetic diversity of the resulting progeny, we performed full-genome amplicon-based deep sequencing on representative replicates of nsP4 variants at residues L383, C483, and W486 in the C483 pocket (**Fig 6A, Fig S6**). Based on the deep sequencing, we called viral variants and quantified their frequency per position in CHIKV genome in each condition. Nonsynonymous mutations are highlighted in gray, synonymous mutations in salmon and mutations in non-coding regions in blue (**Fig 6B-H, Table S1**). In BHK-21 cells (**Fig 6B-D**), as expected, the high-fidelity nsP4 variant C483Y led to a reduced number of minor variants compared to the WT polymerase (**Fig 6C**, quantified in **Fig 7A**). Indeed, C483Y, as a high-fidelity variant, presents a significantly lower mutation frequency, thus directly impacting the subsequent genetic diversity of the new viral populations. In contrast, the nsP4 variant W486Y generated more diversity than nsP4 WT, as we see with the production of a higher number of variants (**Fig 6D**, quantified in **7A**). Interestingly, the nsP4 W486F variant, which reverted in a minority of rescue events, tended to generate less genetic diversity than nsP4 WT (**Fig S7A and B**), suggesting a regulator switch similar to what was shown for C483Y *vs* C483G (18). In mosquito cells, the nsP4 variant C483Y was consistently associated with less genetic diversity than nsP4 WT (**Fig 6F**). Surprisingly, the polymerase variant W486Y replication resulted in a more important genetic diversity in mammalian cells than in mosquito cells (**Fig 6H**). In addition to minor variants, we also quantified the Root Mean Square Deviation (RMSD), and the ratio of nonsynonymous to synonymous mutations (dN/dS ratio) between nsP4 variants and cell types (**Fig 7B, C, F and G**). As a control, we found that the high-fidelity nsP4 C483Y variant showed a lower RMSD in both cell types (**Fig 7B and F**) confirming what has previously been reported (17,18,26). For the nsP4 variants L383F and W486Y, we found slightly increased diversity compared to nsP4 WT, as measured by RMSD for nsP4 W486Y in BHK-21 cells (**Fig 7B**) and nsP4 L383F in C6/36 cells (**Fig 7F**). Regardless, while we do find general trends, none of these were statistically significant.

**Fig 6.**
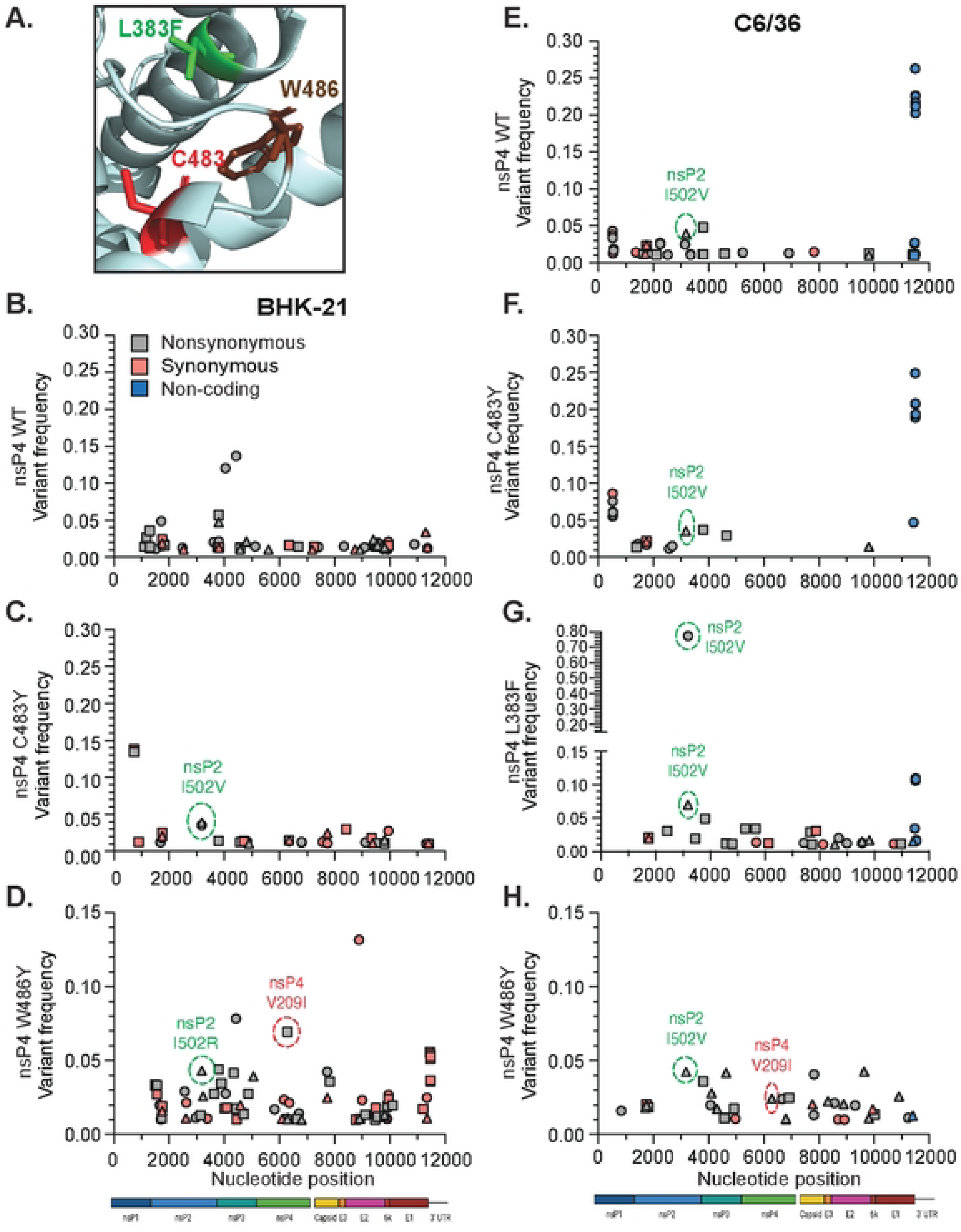
Mutation in the palm subdomain of nsP4 impacts viral genetic diversity. (**A**) Zoom on the C483 pocket in the palm subdomain of nsP4 (PDB: 7Y38) viewed on PyMol. The side chains of the residues of interest are shown. L383F is colored in green, C483 is colored in red, and W486 is colored in brown. Supernatants from three independent biological replicates of the genetically stable variants from the L383, C483 and W486 residues in both cell types (corresponding to the Fig 2) were subjected to deep sequencing. The bioinformatic analysis was performed using a conservative threshold of 1% for the variant calling. Variant frequency by nucleotide position along the genome is mapped for BHK-21 cells (left side) and C6/36 cells (right side). Shown are the minority variant frequencies for BHK-21 cells nsP4 WT (**B**), nsP4 C483Y (**C**), nsP4 W486Y (**D**) and for C6/36 cells nsP4 WT (**E**), nsP4 C483Y (**F**), nsP4 L383F (**G**) and nsP4 W486Y (**H**). Nonsynonymous mutations are colored in grey, synonymous mutations in salmon and non-coding mutations in blue. The variants of interest nsP2 I502V and nsP4 V209I are highlighted in a dashed circle, respectively colored in green or red. A schematic of CHIKV genome is depicted at the bottom of the graphs.

**Fig 7.**
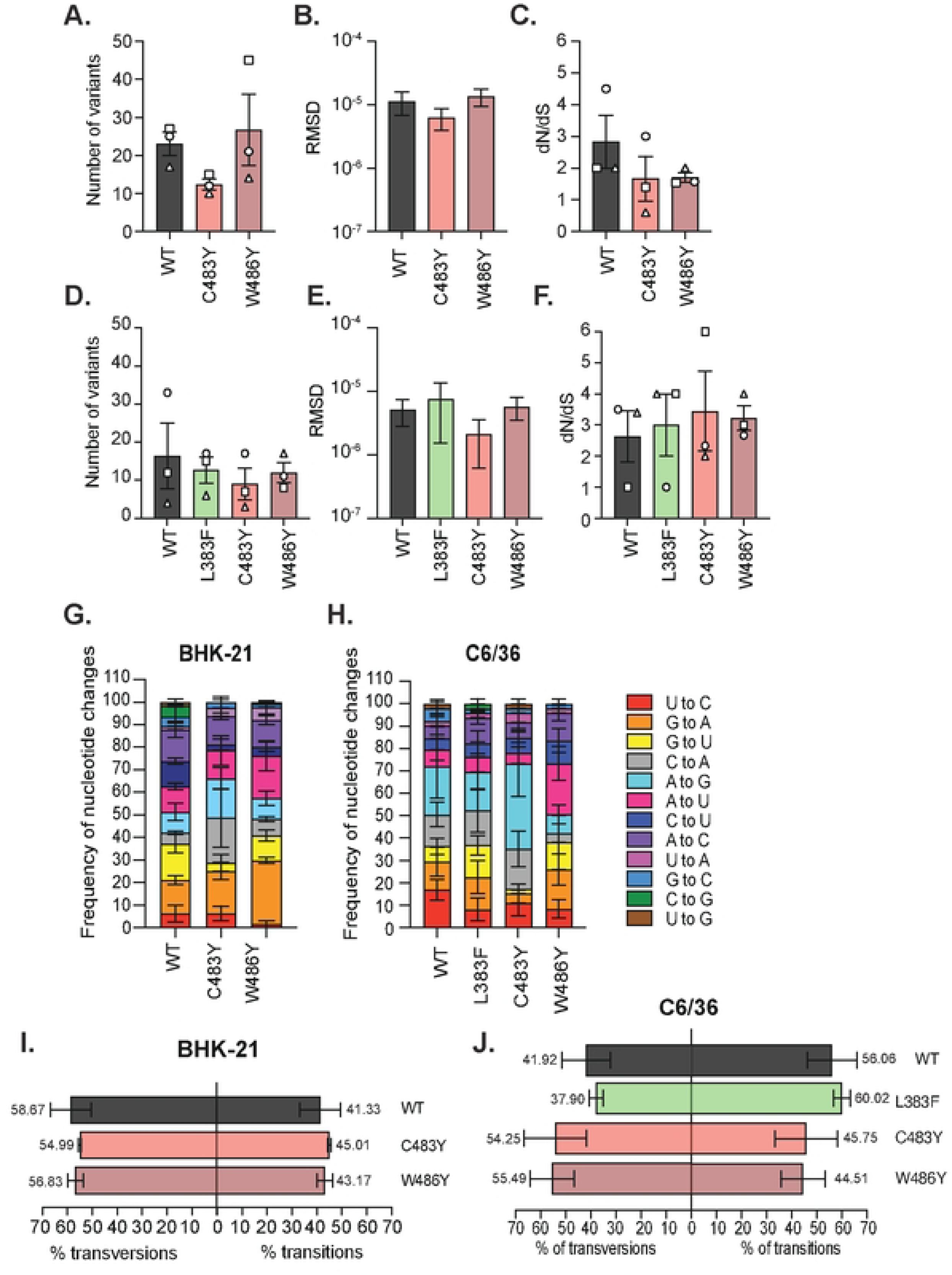
Mutating the palm subdomain alters the type of nucleotide changes made by the polymerase. The BHK-21 cells (**A**) or C6/36 cells (**D**) absolute number of minority variants is presented. The Root Mean Square Deviation (RMSD), a measure of the average diversity generated by the nsP4 variants at each position across the genome, is depicted in (**B**) for the BHK-21 cells and in (**E**) for the C6/36 cells. The dN/dS ratio for each variant in BHK-21 cells (**C**) or C6/36 cells (**F**) is shown. For all panels, a Mann-Whitney *t* test was performed against nsP4 WT but no data reached statistical significance. Graphs show the average and SEM of three biological independent experiments. Analysis of the specific nucleotide changes frequency corresponding to each polymerase variants in BHK-21 cells (**G**) or C6/36 cells (**H**) is presented. Each substitution type is associated to a given color: red for U to C, orange for G to A, yellow for G to U, grey for C to A, light blue for A to G, pink for A to U, dark blue for C to U, purple for A to C, mauve for U to A, medium blue for G to C, green for C to G and brown for U to G. For all panels, a Mann-Whitney *t* test was performed against nsP4 WT but no data reached statistical significance. Graphs show the SEM of three biological independent experiments. The percentage of transitions and transversions events is shown for BHK-21 cells (**I**) and C6/36 (**J**).

In both cell types, we observed the presence across all samples, except for nsP4 WT in BHK-21 cells, of a nsP2 protease domain variant I502V or I502R (**Fig 6, green circle**). Interestingly, the nsP2 I502V variant is strongly selected in the context of the nsP4 L383F variant with a frequency of 77.3% in the first independent biological replicate and 7% in the second (**Fig 6G**). We also observed a second-site mutation in nsP4, V209I (**red circle**), common to the W486Y and F variants (**Fig 6D and H, Fig S7A**). The residue V209 locates in proximity to the residue W486, and both are facing the same nsP2 helicase domain loop (**Fig S7G**), likely compensating for the introduced substitution at position 486 in nsP4. Finally, in C6/36 cells, we observed a significant number of minor variants in the 3’UTR for nsP4 WT and nsP4 C483Y (**Fig 6E-F, blue circles**). However, these variants were not present in the nsP4 L383F, W486Y, or W486F conditions, suggesting that these residues have specific interactions with the CHIKV 3’UTR during replication.

To gain further insight into the molecular biology of the polymerase variants, we analyze the mutation dynamics in the deep-sequenced viral populations and quantified the rate of transition events (substitution of a purine for another one, A to G or G to A substitutions, or a pyrimidine to another one, C to T/U or T/U to C substitutions) and transversion events (substitution of a purine for a pyrimidine or conversely). Even if there are twice as many as transversion combinations possible during replication, transitions are generally favored at a higher frequency because they use a purine and a pyrimidine in the polymerase active site and they tend to generate more synonymous and conservative mutations than transversions, thus limiting their overall impact on viral fitness (27). Importantly, the nsP4 variant L383F in C6/36 cells (**Fig 7H**) presented an increased frequency of transition events (60.02%) while the C483Y and W486Y variant lead to a decreased frequency of transversion events (respectively 45.75% and 44.51%) than the WT polymerase (56.06%) (**Fig 7H**). This phenotype is also observed for the W486F variant in BHK-21 cells (with 63.78% *vs* 41.33% for nsP4 WT) (**Fig S7E**).

In a second time, we analyzed the nucleoside frequency changes in each condition and cell type (**Fig 7I and J**). In mammalian cells, the nsP4 variants C483Y and W486Y presented distinct patterns of nucleoside changes. The nsP4 C483Y variant showed increased A to G (17.38% ± 8.39%) and C to A (19.94% ± 22.57%) substitutions, whereas the W486Y variant generated more G to A (28.33% ± 2.25%) and A to U (18.75% ± 11.51%) substitutions than nsP4 WT (respectively 9.09% ± 6.63% for A to G; 4.92% ± 1.15% for C to A; 14.76% ± 3.46% for G to A and 11.29% ± 2.58% for A to U) (**Fig 7G**). These increases were notably compensated in the case of nsP4 W486Y by a reduced representation of U to C (1.55% ± 2.69%) and U to G substitutions (0.78% ± 1.34%) compared to the WT polymerase (6.25% ± 6.25% and 1.44% ± 2.51% respectively). This result was also seen with the W486F variant which presented a strikingly distinct pattern of nucleoside changes with no U to C, no U to G and no C to G changes, as well as a disproportion of G to A (27.88% ± 4.08% *vs* 14.76% ± 3.46% for nsP4 WT) and C to U (19.87% ± 4.53% *vs* 11.23% ± 2.20% for nsP4 WT) substitutions (**Fig S7F**). In mosquito cells, the variants C483Y, L383F, W486Y also displayed altered nucleoside change frequencies compared to the WT polymerase (**Fig 7J**). In particular, all nsP4 variants produced few U to C changes (16.92% ± 7.96% for nsP4 WT *vs* 8.10% ± 8.91% for nsP4 L383F, 16.51% ± 3.16% for nsP4 C483Y and 8.33% ± 7.21% for nsP4 W486Y), while nsP4 C483Y produced more A to G substitutions (23.66% ± 6.94%) and W486Y again favored A to U changes (22.73% ± 13.11%), as in mammalian cells, compared to nsP4 WT (respectively 21.46% ± 24.74% and 7.58% ± 8.44%).

These changes in the frequency of substitution events show that, in addition to affect the proper viral replication and genetic diversity of the progeny, introducing mutations in the polymerase strongly affect the error bias of nsP4.

Taken together, these results suggest an alteration of the NTP discrimination and/or binding by the polymerase variants L833F, W486F and W486Y, affecting the nature of the errors introduced by the viral polymerase upon replication. Finally, the changes in viral diversity observed in the nsP4 variants have strong implications for the viral progeny and may explain, at least in part, the alterations of viral growth.

## Discussion

Alphaviruses are significant human health threats, yet there are no or limited therapeutic strategies available. One potential broad-spectrum antiviral target is the highly conserved RNA-dependent RNA polymerase, highlighting the need to understand how the alphavirus polymerase functions. In this study, we took advantage of the Coxsackievirus B3 (CVB3) 3 polymerase, a well-characterized model for positive-sense RNA virus polymerase study, and the predictive power of homology-modeling to unravel unknown key CHIKV polymerase activity and fidelity determinants. With this objective, we predicted a set of variants from seven conserved CHIKV nsP4 residues based on the CVB3 3D^pol^ structure. We hypothesize they may play critical roles in nsP4 replication and function and in the changes taking place within the polymerase during active site closure and catalysis.

When we tried to rescue each nsP4 variant in mammalian cells, we found that only the variants C483Y, W486L and W486Y remained genetically stable. As previously established (17,18), nsP4 C483Y is a high-fidelity variant of the polymerase and as such it is expected that it stays genetically stable. Given the stability of nsP4 W486Y, we hypothesize that this nsP4 variant is associated with either an increased or a similar fidelity to nsP4 WT and/or that the substitution with a tryptophan at position 486 of the polymerase is relatively well tolerated by the polymerase. Interestingly, CHIKV nsP4 variants I312V and L368F were not genetically stable in mammalian cells, whereas the corresponding CVB3 3D pol variants were stable in human cells (22). More specifically, in CVB3, these variants were associated with a slighter or a significantly higher mutation rate compared to the WT, even if they were still replicating with the same efficiency as the WT polymerase (22).

On the contrary, in mosquito cells, all rescued nsP4 variants with the exception of the nsP4 I312V variant were genetically stable. The I312 residue is located close to the polymerase ring finger (**Fig 1A and Fig 8**). The finger subdomains are highly flexible subdomains involved in conformational changes during polymerization that maintain the active closed structure of the catalytic site and are required for RNA template binding (9,28). Given that the I312V variant was unstable in both mammalian and mosquito cells, it suggests a potential key role for the ring finger in both species. Regardless, the increase in genetic stability over mammalian cells is interesting as it has been shown that the mosquito host leads to a strong purifying selection on viral populations as a consequence of a series of consecutive severe biological bottlenecks (29).

**Fig 8.**
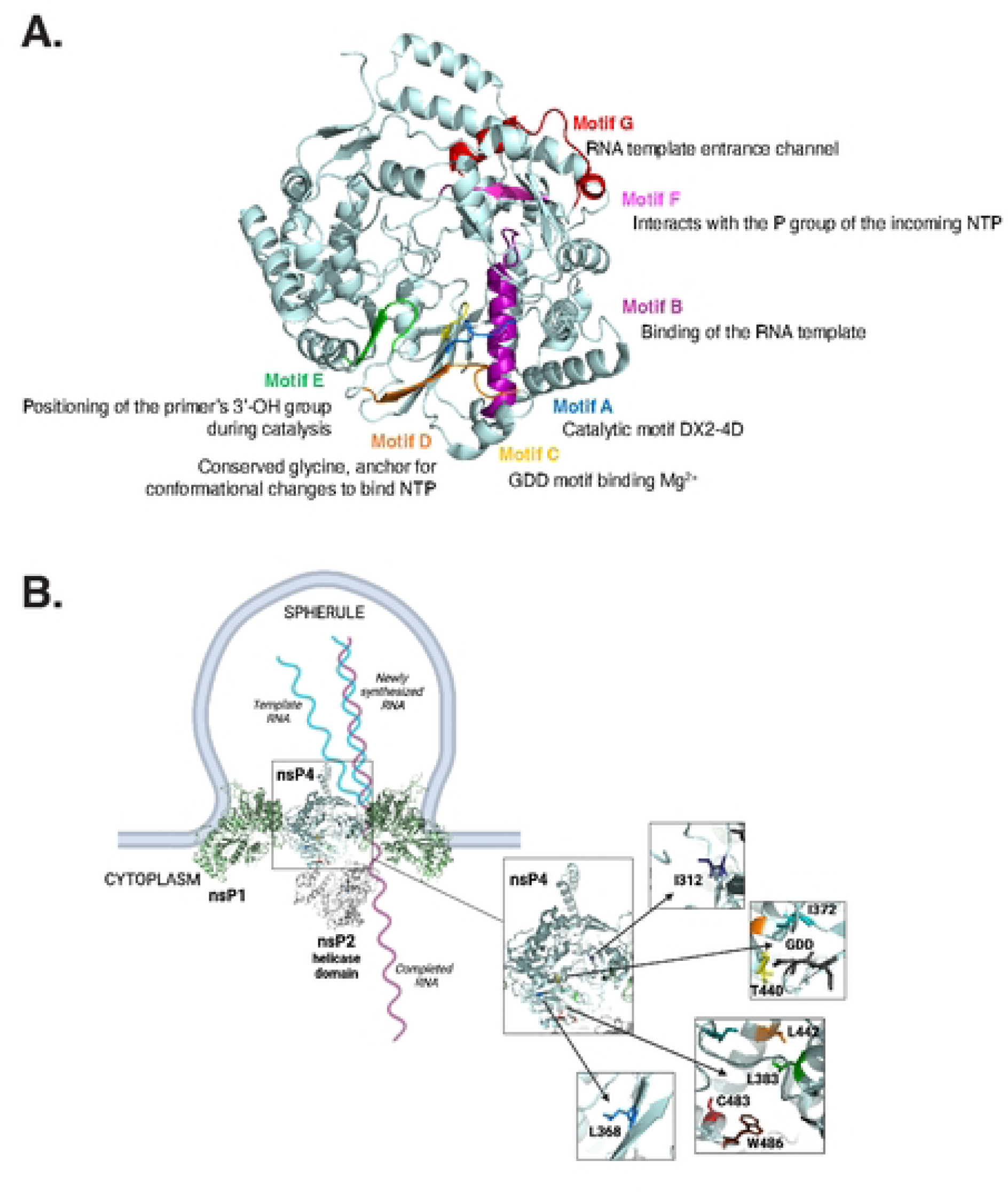
nsP4 variants are close to crucial conserved enzymatic activity motifs and the helicase nsP2 in the context of the core replicase complex. **(A)** nsP4 3D structure (PDB: 7Y38) showing in color each of the conserved positive-sense RNA-dependent RNA polymerase motifs and their respective function. Motif A is depicted in blue, motif B in purple, motif C in yellow, motif D in orange, motif E in green, motif F in pink, and motif G in red. (**B**) nsP4 in the context of the core replicase complex (PDB: 7Y38). A zoom on nsP4 and the different pockets corresponding to the nsP4 variants is shown. The residues L383, C483 and W486 are located at the interface between nsP4 and nsP2. Schematic made with Biorender.

Strikingly, in both cell lines, no nsP4 variant from the residue I372 and T440 could be rescued, nor did they produced intracellular RNA or viral proteins, indicating a strict requirement for these polymerase residues in RNA synthesis. I372 is in the conserved motif A and in immediate proximity to the catalytic aspartic acid residue at position 371 involved in the coordination of the magnesium cations during catalysis (**Fig 1B**, **Fig 8A and B**) (9, 6, 20). This observation underlines the strong evolutionary constraints taking place in close vicinity to the polymerase active site, an idea supported by the Protein Contact Atlas predicted interactions that indicates strong atomic level interactions between the I372 residue and the catalytic residues D371 and D466 (data not shown) (30). Importantly, motif A has been described to align with motif C upon binding of the correct dNTP, initiating the closed and active RdRp conformation required for polymerization (28), thus emphasizing the pressure on this specific region of the protein (**Fig 8A**).

Interestingly, in mosquito cells, variants of nsP4 residues L368, L383, and L442 produce extracellular and intracellular viral RNA and proteins, yet do not produce infectious virus particles, suggesting a role for this region of nsP4 in particle assembly. While we did not find viral glycoproteins in the supernatant for these variants, we confirmed the presence of intercellular extensions for nsP4 WT, as being established as actin/tubulin positive and of more than 10 μm in length (24). These extensions were also harbored by nsP4 variants L368F and L368V that showed WT-like structural protein expression patterns. In contrast, the variants L383A and L442A, despite expressing structural proteins, displayed altered actin distribution. This observation is of particular interest as it has been shown that HIV (31,32), Influenza A virus (33) and flaviviruses (34) subvert the actin network at their own advantage in order to redirect viral components towards the plasma membrane for viral assembly, although it seems that it could be negatively affecting the step of budding itself (35).

Residue L368 is interesting as it is located just before the conserved motif A in the palm subdomain but is also quite structurally isolated from both the catalytic site pocket and the C483 pocket (**Fig 8B**). In addition, residue L442 is located inside motif B, involved in the binding of the RNA template (28), allowing us to speculate that even a slight change in amino acid side chain length, as in between a leucine and an alanine, has a dramatic impact on the correct positioning of the template in the polymerase. Interestingly, the L383 residue directly faces the L442 residue (**Fig 8B**). Importantly, networks of isoleucine, leucine and valine residues (ILV clusters) have been described as crucial for stability in high-energy partially folded protein states (36), that are likely observed in RdRp upon catalysis, but hard to capture by experimental biology (37).

Another explanation to the non-rescue of several nsP4 variants in mosquito cells was either a delay in viral production and/or the generation of non-infectious viral particles, also known as defective viral genomes (DVGs). DVGs are sub-viral particles that generally contain truncations of the viral genome, rendering it unable to undergo a full replication cycle except with the presence of a helper virus, able to complement the lost viral function (38). When these particles are interfering with viral replication, they are named defective interfering particles (DIPs). A study by Poirier *et al.*, has shown using the low-fidelity SINV polymerase C482G, the analog of the C483G variant for CHIKV nsP4, that by recombining at higher rates than the WT, the polymerase produces large amounts of DIPs (39). In our hands, CHIKV nsP4 C483G was not able to be well rescued in mosquito cells contrary to mammalian cells, likely via lethal mutagenesis, this low-fidelity polymerase incorporating too many errors in the genome for it to be viable.

Nonetheless, the idea of an nsP4 role in assembly is intriguing. Putting back the nsP4 variants studied in this work in the context of the replication complex solved by Tan *et al* (**Fig 8B**), residues L368, L383, and L442 locate to the cytoplasmic side of the spherule near the interface of the helicase domain of nsP2 with nsP4. While this structure lacks other viral proteins including nsP3, we could hypothesize that these sites also contain the viral capsid and glycoproteins to facilitate assembly. Indeed, a likely proximity between viral replication complexes and viral assembly sites has been evoked (40,41) and could facilitate packaging of genomic RNA. Interestingly, genomic RNA is selectively packaged in the nucleocapsid whereas present small amounts in the cell during replication compared to the subgenomic RNA. It has been described that viral packaging requires both RNA-protein interactions and protein-protein interactions involving respectively, and chronologically, genomic RNA-capsid, capsid/capsid then capsid/E2 interactions (42). Moreover, work with Venezuelan Equine Encephalitis virus (VEEV) and Semliki Forest virus (SFV) has suggested that nsP2 could be involved in this presentation, as it contains packaging signal sequences (43) and seems to regulate viral assembly of infectious virions and tightly interacts with the helix I of capsid protein (44). In addition, nsP2 has been shown to interact genetically and physically with nsP4 (13,19,45). Finally, intrinsically disordered domains, through the high flexibility they confer to proteins, are interesting candidates that could mediate these suggested interactions. Both capsid and nsP4 N-terminal domains are intrinsically disordered regions, respectively already demonstrated to play roles in gRNA binding during assembly (42,46) and active polymerization (9,47). Although our study did not provide experimental evidence of nsP4 being part of the viral assembly complex, further work is needed to investigate this plausible polymerase role in the late replication steps.

Finally, as RdRps are replicating the genome and intrinsically incorporating errors at the same time, they are generating important genetic diversity. Except for the coronaviruses, most RNA viruses do not possess a proof-reading activity (48,49). This is why we wondered if mutations in the palm domain of the polymerase, especially those in close proximity to the C483 residue, a universal alphaviruses fidelity determinant, will affect the genetic diversity of the progeny. To this end, using deep sequencing of the progeny of variants at residues L383, C483, and W486 in both cell types, we observed a reduced genetic diversity for the high-fidelity variant C483Y in both cell types, which was expected due to the low mutation frequency of this variant (17). Interestingly, when we looked at the variant frequency across the genome in each condition, we noted the presence of a second-site mutation in nsP4 (V209I) in both mammalian and mosquito cells for the nsP4 W486 variants. The nsP4 residue V209 is located close to the residue W486, and both residues are facing the same nsP2 helicase domain loop. Importantly, as the viral protease (11) as well as the helicase (12), unwinding the RNA strands during replication, nsP2 is a component of the core replicase complex and works with nsP4 to control replication fidelity (13). nsP2 sits on the ring of nsP1 molecules on the cytoplasmic side of the replication complex and is facing nsP4, itself buried in the spherule (19). Interestingly the substitution between the original valine and an isoleucine at the position 209 of nsP4 is shortening the distance between nsP4 and this helicase domain nsP2 loop, thus likely reinforcing their interaction. Another mutation of interest happened to be in nsP2 (I502), again in both cell types but this time associated to all polymerase variants, including the nsP4 WT in mosquito cells. Strikingly, this substitution at position 502 between the original isoleucine and a valine or an arginine (**Fig 6C-H**), appears to be strongly selected in the nsP4 variant L383F in the context of mosquito cells. The I502 residue is located in the protease domain of nsP2 that also hosts residues associated with several other minor variants in nsP2, including the variants L708P. To conclude, these observations emphasize the previously characterized functional link between nsP4, the RdRp, and nsP2, the viral helicase and protease of CHIKV.

While our work describes important domains for CHIKV nsP4 function, genetic stability, replication through alternate hosts, and particle assembly, it possesses several limitations. First, the cell lines we selected for this study are particularly permissive to CHIKV infection and more broadly to arboviral infections. Indeed, BHK-21 cells are defective at producing interferon (50) and C6/36 cells do not have a competent RNAi pathway due to a defect in Dicer-2 (51,52). Thus, looking at nsP4 variants replication in RNAi- or interferon-competent cells, like U4.4 cells, also derived from *Aedes albopictus* mosquitoes, or human fibroblasts will give us insight into the impact of this immune selective pressure on viral replication itself. In addition, temperature could play a major role on the replication rate and virus stability. As BHK-21 and C6/36 cells grow at 37°C and 28°C respectively, future studies addressing the role of temperature will be important to understand nsP4 biology. Finally, via our targeted approach of designing a specific panel of mutants, we only have a partial view of functional residues in the polymerase. Expanding outside of the palm domain and ring finger will be key for understanding how nsP4 functions.

In conclusion, these studies investigate how discrete domains of the CHIKV polymerase function in the complete alphavirus life cycle. We identified novel determinants required for polymerase genetic stability and found a previously unknown role for nsP4 in viral assembly. Furthermore, it underlines the differential selective pressures encountered by the polymerase during infection. Importantly, we demonstrated that the palm domain of the polymerase plays a crucial role in the generation of subsequent viral genetic diversity, likely via its role in binding and/or discriminating NTPs during RNA synthesis. Future work is now required using *in vivo* models for a more integrated and comprehensive overview, and, in particular, to provide insight on the attenuation of these viruses in their natural environment giving the alteration of the resulting genetic diversity observed *in vitro*. As we highlighted important host-specific pressures, it would be interesting to see if our observations on CHIKV nsP4 could be expanded to other alphaviruses, notably to alphaviruses known to circulate in *Culex spp.* species (SINV, RRV or VEEV) or to have birds as a natural reservoir (SINV, RRV).

## Materials and Methods

### Cell lines

Baby hamster kidney cells (BHK-21 CCL-10, ATCC) were grown in Dulbecco’s Modified Eagle Medium (DMEM, Corning, #10-017-CV) supplemented with 10% fetal bovine serum (FBS, Atlanta Biologicals), 1% N-2-hydroxyethylpiperazine-N’-2-ethanesulfonic acid (HEPES, Gibco, #15630080) and 1% MEM nonessential amino acids (MEM-NEAA, Gibco, #11140050). *Aedes albopictus* cell clone C6/36 (CRL-1660, ATCC) was grown in Leibovitz’s medium (L-15, Corning, #10-045-CV) supplemented with 10% FBS, 1% tryptose phosphate broth (Gibco, #18050039) and 1% MEM-NEEA, Gibco, # 11140050). Vero cells (CCL-81, ATCC) were grown in DMEM supplemented with 10% newborn calf serum (NBCS, Sigma). All the cell lines were maintained at 37°C under controlled CO_2_ atmosphere (5%) with the exception of C6/36 maintained at 28°C. Each cell line was tested monthly for mycoplasma using the Lookout Mycoplasma PCR detection kit (Sigma-Aldrich, # MP0035) and confirmed mycoplasma free. BHK-21 and C6/36 cells of passage 10 or less were used for every experiment (or alternatively less than 1-month old, maintained cells).

### Homology modeling

An initial model for the CHIKV nsp4 structure was developed by a combination of homology modeling and threading onto the Coxsackieviurs B3 polymerase structure (PDB: 3DDK), using the programs I-TASSER (53), Swiss-model (54), and Phyre (55). This approach provided a reasonable model for the conserved active site motifs in the palm domain that allowed us to identify residues likely to generate fidelity-modulating variants based on our prior work with coxsackievirus B3 polymerase (23). Residues were chosen based on their proximity to the pre-existing C483Y variant (17) or having locations likely to impact the palm domain-based movement of motif A that underlies the active site closure mechanism of positive-strand RNA virus polymerases (28, 56). The locations of these residues were later validated when the structures of Sindbis and Ross River virus and ONNV (12) polymerases were solved.

### Molecular biology and plasmid mutagenesis

To generate wild-type (WT) chikungunya virus (CHIKV) (Strain 06-049; AM258994) and nsP4 variants, a modified chikungunya virus (CHIKV) infectious clone containing an AvrII restriction site at the 3’ end of the subgenomic promoter was used (pCHIK-FLIC-AvrII) (57). The nsP4 variants were introduced into a small cloning vector via site-directed mutagenesis using the primers in Table 2 and Sanger sequenced to confirm the variant. The AgeI/AvrII restriction fragment was then subcloned from each nsP4 variant into pCHIK-FLIC-AvrII and each plasmid was Sanger sequenced to confirm the nsP4 variants.

**Table 2:**
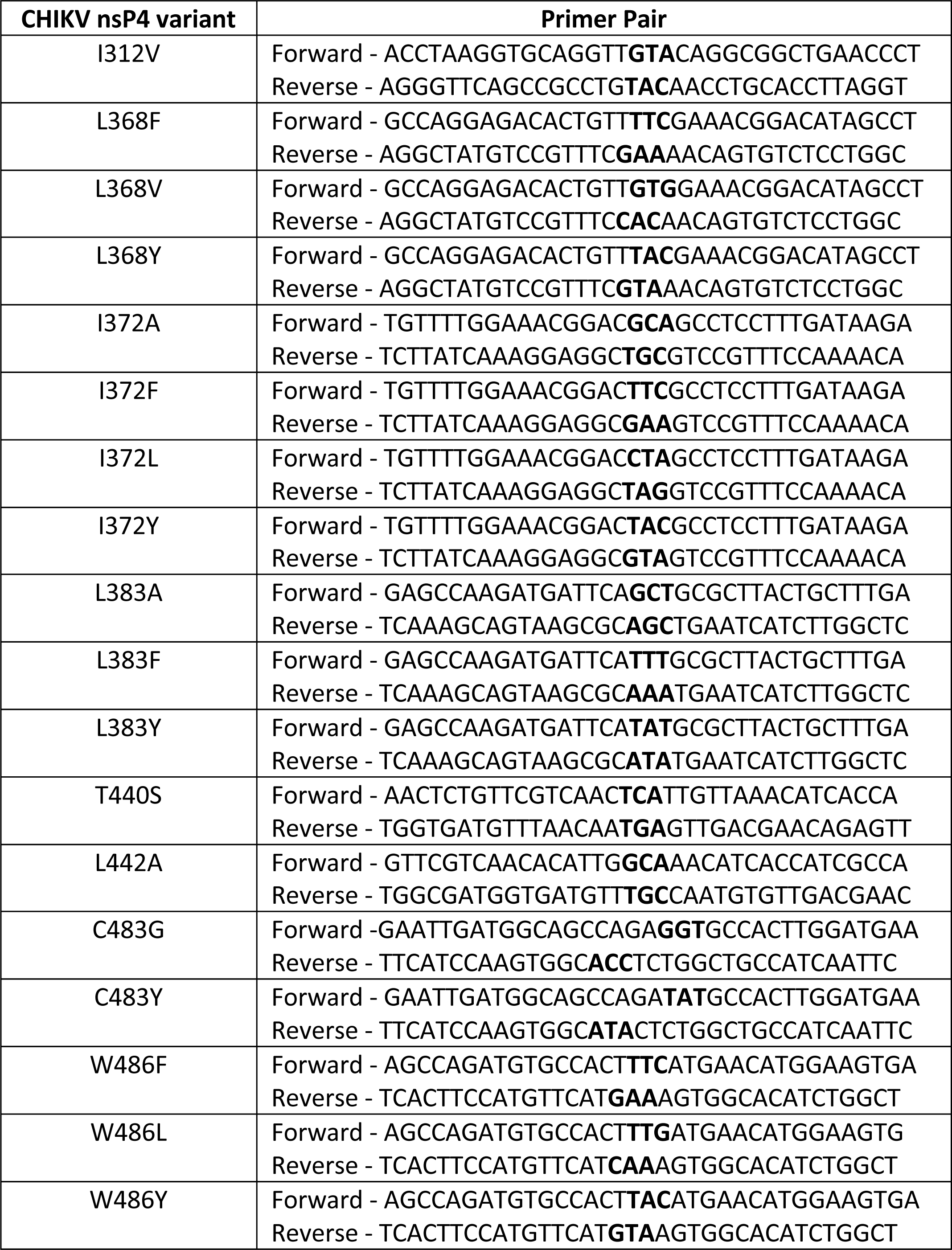
Mutagenesis primers used in this study.

To generate virus, full-length WT CHIKV and nsP4 variants were generated by linearization of 10 μg of plasmid overnight with NotI restriction enzyme. The next day, the linearized product was purified by phenol:chloroform extraction and ethanol precipitation and re-suspended in RNAse free water. The purified product was then *in vitro* transcribed using the mMessage mMachine SP6 kit (Invitrogen, # AM1340) following manufacturer’s instructions. *In vitro* transcribed RNA was purified by phenol:chloroform extraction and ethanol precipitation and finally re-suspended in RNAse free water. After Nanodrop 2000 Spectrophotometer (ThermoScientific) quality control and concentration measurement, every RNA was diluted to 1 μg/μL and aliquoted before storage at −80°C.

### Biosafety

All experiments with WT CHIKV and CHIKV nsP4 variants were performed in a biosafety level 3 (BSL3) laboratory at NYU Grossman School of Medicine (New York, NY USA).

### Plaque assay

Viral infectious particle progeny was quantified by titration on Vero cells in 12-well plates. Briefly, Vero cells were seeded at a concentration of 350,000 cells/well. The next day, ten-fold dilutions of each supernatant were made in DMEM and 200 μL of virus was added to each well. Cells were incubated during 1 hr at 37°C. After virus adsorption, 1.5 mL of pre-warmed plaquing medium made of DMEM-2% NBCS-1% Antibiotic-Antimycotic 100X (Invitrogen, # 15240062) was added per well. Plates were incubated for 3 days at 37°C. Plates were then fixed with 1.5 mL of 4% formalin per well (Fisher Scientific, # SF1004) for 1 hr at room temperature. After fixation, formalin and agarose plugs were removed, followed by crystal violet (Sigma-Aldrich, # HT90132) staining for 15 mins. Finally, plates were washed in diluted bleach (10% Clorox bleach, Fisher Scientific, # 50371500) and left open 24 hours to dry at room temperature. The number of plaques by dilution where plaques were observed in each condition was then counted and the number of infectious viral particles per initial inoculum determined.

### RNA extraction and RT-qPCR

For extracellular RNA quantification, RNA extraction was performed using the PureLink RNA mini kit (Invitrogen, # 12183025). Briefly, samples in Trizol were incubated at room temperature for 5 mins. Then, 0.2 mL of chloroform per 1 mL of Trizol reagent used was added and the tube was vigorously shaken for 15 secs, followed by 3 mins incubation at room temperature. Samples were then centrifuged at 12,000 x g for 15 mins at 4°C. The colorless upper phase was then transferred onto a new tube and mixed with an equal volume of 75% ethanol. Samples were then vortexed and transferred onto a column. Several washes with the kit buffers were then performed according to manufacturer’s instructions. Finally, after drying of the membrane, 100 μL of RNAse-water was added on each column to elute extracellular RNA before storage at −80°C.

For intracellular RNA quantification, RNA extraction was performed using Trizol reagent (Invitrogen, # 15596026). Briefly, after cell lysis into Trizol, 1/5^th^ of the total volume of chloroform was added in each sample and vortexed for 10 secs, followed by 5 mins incubation at room temperature. Samples were then centrifuged at 12,000 x g for 15 mins at 4°C to allow for phase separation. Following the spin, the upper aqueous phase was transferred to a new tube containing 500 μL of molecular grade isopropanol (Absolute isopropanol molecular grade, Fisher Scientific, # 67-63-0) and 1 μL of Glycoblue co-precipitant (Invitrogen, # AM9515). RNA was precipitated overnight at −20°C. The next day, the samples were vortexed and then centrifuged at 12,000 x g for 20 mins at 4°C. Supernatant was removed, and the RNA pellet was washed two times with 700 μL of 75% molecular grade ethanol (Absolute ethanol molecular grade, Fisher Scientific, # BP28184). Finally, ethanol was completely removed, and the pellet was resuspended in 100 μL of RNAse free water. Samples were normalized to 100 ng/μl with RNAse free water. RNA quality and concentration was then checked on a Nanodrop 2000 Spectrophotometer (ThermoScientific) before storing samples at −80°C.

For RT-qPCR, the number of viral genomes/mL was quantified in a transparent 96-well plate by TaqMan RT-qPCR on the previously extracted RNA (see above). To do so, the TaqMan RNA-to-Ct One-Step-RT-PCR kit (Applied Biosystems, # 4392938) was used along with CHIKV primers and a FAM-probe targeting the nonstructural protein 4 (CHIKV Fwd IOLqPCR-6856F: TCACTCCCTGCTGGACTTGATAGA / CHIKV Rev IOLqPCR-6981R: TTGACGAACAGAGTTAGGAACATACC / CHIKV probe IOLqPCR6919-FAM: AGGTACGCGCTTCAAGTTCGGCG). The following mix was done for each reaction: 12.5 μL of Taqman RT-qPCR mix 2X, 6.25 μL of RNAse-free water, 0.25 μL of CHIKV Fwd IOLqPCR-6856F, 0.25 μL of CHIKV Rev IOLqPCR-6981R, 0.15 μL of CHIKV probe IOLqPCR6919-FAM, 0.63 μL of Taqman enzyme mix and 5 μL of sample. A standard curve was generated for each plate using ten-fold dilutions of *in vitro* transcribed CHIKV RNA. Each experimental sample and standard curve dilution was done in technical duplicates. The plate was then sealed, centrifuged briefly and ran on a QuantiStudio 3 real-time machine (Applied Biosystems). The amount of CHIKV RNA molecules per mL of sample was determined via the associated standard curve.

### Virus replication kinetics

BHK-21 and C6/36 cells were seeded in a 24-well plate at a respective concentration of 75,000 cells/well or 150,000 cells/well to be 70-80% confluent the next day for transfection. 24 hours later, a total of 0.5 μg of RNA per well was transfected with Lipofectamine MessengerMAX Transfection Reagent (Invitrogen, # LMRNA001) at a respective ratio of 1:2 (RNA:Lipofectamine). After liposome formation, each mix was added dropwise on top of the corresponding wells and cell culture medium was left according to manufacturer’s instructions. The next day, all the supernatant was removed and kept for plaque assay. Cells were then washed 3 times with 1X phosphate buffered saline (PBS) and fresh complete cell medium was added back. Plates were then incubated at the corresponding temperature according to the cell type. At 48 hpt, 60 μL of supernatant was taken and kept for plaque assay before to add back 60 μL of fresh complete media and incubate back the plates. Finally, at 72 hpt, all the supernatant was removed and kept for plaque assay and extracellular RNA extraction.

### Immunofluorescence staining and confocal microscopy

BHK-21 and C6/36 cells were seeded in a 24-well plate on 12 mm round coverslips (Round German coverslip 12 mm, Bellco Glass, # 1943-10012A) at a respective concentration of 75,000 cells/well or 150,000 cells/well to be 70-80% confluent the next day for transfection. 24 hours later, cells were transfected with 0.5 μg of *in vitro* transcribed RNA as described above. 24 hours later, the wells were fixed by adding on top of the supernatant 10% formaldehyde (Formaldehyde Aqueous Solution EM grade, 32%, Liquid Electron Microscopy Sciences, # 15700) at a volume-to-volume ratio for 1 hour at room temperature. After fixation, the supernatant was removed, and cells were washed 3 times with Phosphate-Buffered Salt solution 1X (Corning, ref: 21031CV) before proceeding to the immunofluorescence staining.

Free aldehyde groups from formaldehyde were neutralized with 50 mM ammonium chloride (NH_4_Cl) for 10 mins at room temperature. After several PBS washes, cells were permeabilized 0.1% Triton X100 (Triton X-100, Fisher BioReagents, # BP151100) for 15 mins and then blocked with 2% bovine serum albumin (BSA, Millipore Sigma, # H2519) for 30 mins at room temperature. Cells were then incubated via inversion of the coverslip on 30 μL of primary antibodies diluted at 1:300 in the 2% BSA solution, for 1 hour at room temperature in a wet chamber. After incubation, coverslips were put back in the plate, washed several times with PBS and then incubated with fluorophore-conjugated secondary antibodies, diluted at 1:1000, for 45 mins. The following primary antibodies were used: anti-CHIKV nsP4 (rabbit, a gift provided by Dr. Andres Merits at the University of Tartu, Estonia), anti-CHIKV E2 (mouse, the following reagent was obtained through BEI Resources, NIAID, NIH: Monoclonal Anti-Chikungunya Virus E2 Envelope Glycoprotein, Clone CHIK-263 (produced *in vitro*), NR-44003), anti-CHIKV Capsid (rabbit, CHIK-122; provided by Dr. Andres Merits) and anti-Phalloidin-A488 (F-actin, ActinGreen 488 Ready Probes reagent, Invitrogen, # R37110). Coupled antibodies were always incubated after the non-coupled ones, if used in the staining. As secondary antibody, either a goat anti-rabbit IgG-Alexa 488 (Invitrogen, # A-11008), a donkey anti-mouse IgG-Alexa 488 (Invitrogen, # A-31570), a goat anti-rabbit Alexa 647 (Invitrogen, # A-21244) or a donkey anti-mouse Alexa 555 (Invitrogen, # A-31570) were used. Finally, cells were labeled with DAPI (Invitrogen, # D3571) and coverslips mounted in ProLong Diamond Antifade (Molecular Probes, ref: P36965) on slides (SuperFrost End Slide 1 mm thick, Electron Microscopy Sciences, # 71867-01). After drying, coverslips were sealed using transparent nail polish. Slides were then stored at −80°C until acquisition on a Zeiss LSM880 confocal microscope (Carl Zeiss). Images were finally analyzed with the Fiji (Image J) software.

### Western Blotting

To analyze intracellular proteins, BHK-21 and C6/36 cells were seeded in duplicate in a 6-well plate at a respective concentration of 300,000 cells/well or 600,000 cells/well to be 70-80% confluent the next day for transfection. 24 hours later, a total of 2.5 μg of RNA per well was transfected with Lipofectamine MessengerMAX Transfection Reagent (Invitrogen, # LMRNA001) at a respective ratio of 1:2 (RNA:Lipofectamine). After liposomes formation, each mix was added dropwise on top of the corresponding wells and cell culture medium was left 24 hours before being removed and replaced with fresh complete media until the end of the experiment. Both supernatant and cells were harvested 48 hours post-transfection to perform plaque assay, intra- and extra-cellular RT-qPCR, Western Blotting and deep-sequencing.

For western blotting, the wells of each 6-well plate were scrapped on ice in PBS after keeping the supernatant for plaque assay and RT-qPCR on viral progeny. Cells were then centrifuged at 5,000 rpm for 5 mins at 4°C. PBS was removed and 60 μL of lysis buffer (1X TBS, 1 mM EDTA, 1% Triton X-100, 1% protease inhibitor) added into each tube. After 1 hour of incubation at 4°C, 60 μL of 2x Laemmli buffer with 10% 2-®-mercaptoethanol was added per tube for a final volume of 120 μL. Lysates were then heat-denatured for 10 mins at 95°C and spun for 5 mins at 10,000 x g before storage at −20°C. For SDS-PAGE, samples were thawed at room temperature and 10 μL of sample per well was loaded in a 15-well gel of a given acrylamide concentration depending on the protein blotted subsequently (10% acrylamide for nsPs and E2, 12% acrylamide for Capsid). After 50 mins of migration, the proteins on the gels were transferred onto an activated polyvinylidene difluoride (PVDF) membrane (Immobilon from Millipore, # IPFL00005) and then blocked with 1X TBS-0.1% Tween-20-5% dry milk for 1 hour at room temperature on a rocker. The membrane was then transferred into the corresponding primary antibody (1:5000 dilution) and incubated during 1 hour at room temperature or overnight at 4°C. Blots were incubated with the following primary antibodies: anti-CHIKV nsP4 (rabbit, provided by Dr. Andres Merits), anti-CHIKV E1 (rabbit, a gift provided by Dr. Gorben Pijlman at the University of Wageningen, Netherlands), anti-CHIKV E2 (mouse, CHIK-48; a gift provided by Dr. Michael Diamond at Washington University at St. Louis, USA), anti-CHIKV Capsid (rabbit, CHIK-122), and actin (mouse monoclonal, Invitrogen, # ACTN05 C4). Three washes for 5 mins with 1X TBS-Tween-20 were performed and the membranes were finally incubated with the corresponding secondary antibody (1:10000 dilution). Depending on the host species of the primary antibody, either a goat anti-mouse-HRP antibody (Invitrogen, ref: 31430) or an anti-rabbit IgG-HRP antibody (Invitrogen, ref: 31460) was used. After another round of washes, the membrane was subjected to revelation via addition of chemiluminescent substrate reagents (Pierce SuperSignal WestPico Plus, ThermoScientific, ref: 34580) at a ratio of 1:1. Both a chemiluminescent and a colorimetric image were taken for each membrane and antibody revelation. Subsequently, membranes were stained with Coomassie blue and dried at room temperature to keep a trace of the protein content of each sample. Images were analyzed using ImageLab (version 6.0.1).

### Purification of supernatants for western blot on extracellular proteins

To isolate proteins from the supernatants, Amicon columns were used (Amicon Ultra-0.5 Centrifugal Filter Units, cut-off of 10,000 kDa, Millipore Sigma, ref: UFC501024). After thawing the supernatants, 500 μL of each sample was added on each column and then spun for 15 mins at 14,000 x g at room temperature according to manufacturer’s instructions. The flow-through was then discarded and each column inverted on a new tube to be reverse spun down for 15 mins at 1,000 x g at room temperature. A final volume of 20 μL purified proteins was recovered, and the proteins were transferred to a new tube before adding 50 μL of lysis buffer. Samples were incubated 1 hour at 4°C and then 50 μL of 2x Laemmli with 10% ®-mercaptoethanol. Finally, each sample was heat denatured for 10 mins at 95°C and then frozen down at −20°C until SDS-PAGE as described above.

### CHIKV full genome PCR and Sanger sequencing

For Sanger sequencing, the purified extracellular RNA was first used to generate cDNA (Maxima H Minus First Strand cDNA Synthesis Kit, Thermo Scientific, ref: K1652). 1 μL of random primers and 1 μL of dNTPs were mixed with 13 μL of extracted RNA per reaction in a PCR tube. Samples were first subjected to a PCR step at 65°C for 5 mins then chilled on ice and spun before adding a mix of 4 μL of RT buffer and 1 μL of reverse transcriptase enzyme (Maxima H RT) per reaction. The samples were then put at 25°C for 10 mins to allow annealing of the random hexamer primers, followed by 30 mins at 50°C for synthesis of the cDNA. Finally, the reaction is terminated by heating at 85°C for 5 mins. Generated cDNA is then stored at −20°C until use. To generate CHIKV amplicons, all cDNA reactions were subjected to a specific PCR reaction for CHIKV genome amplification with the Phusion PCR kit (Phusion High-Fidelity DNA polymerase kit, Thermo Scientific, ref: F530L). 5 pairs of primers were used to span all CHIKV genome (Table 3, CHIKV PCR primers), namely F1 / F2 / F2.5 / F3 and F4. A 50 μL reaction was made for each sample as follow: 10 μL of 5X Phusion HF buffer, 1 μL of 10 mM dNTPs, 5 μL of 10 μM forward primer, 5 μL of 10 μM reverse primer, 0.5 μL of Phusion DNA polymerase and 4 μL of cDNA. The following PCR program was then applied: an initial denaturation at 98°C for 30 secs, then 35 cycles of a denaturation step at 98°C for 10 secs, annealing of the primers at 51°C for 30 secs and extension of the product at 72°C for 2 mins. A final extension at 72°C for 10 mins was then performed followed by a cool down at 4°C. Amplification of the fragments was then checked on a 1% agarose gel run at 120 volts for 35 mins. All positive samples were purified with the NucleoSpin Gel and PCR Clean-up kit by Macherey-Nagel (ref: 740609.50S). Briefly, DNA was bound onto the column and spun at 11,000 x g for 30 secs. Flow-through was discarded and the column was washed 2 times before drying with a spin at 11,000 x g during 1 min. DNA was then eluted and quantified on a Nanodrop 2000 Spectrophotometer (ThermoScientific). Purified samples were Sanger sequenced at Genewiz (Azenta) using a specific set of primers (**Table 3**, CHIKV Sanger sequencing primers).

**Table 3:**
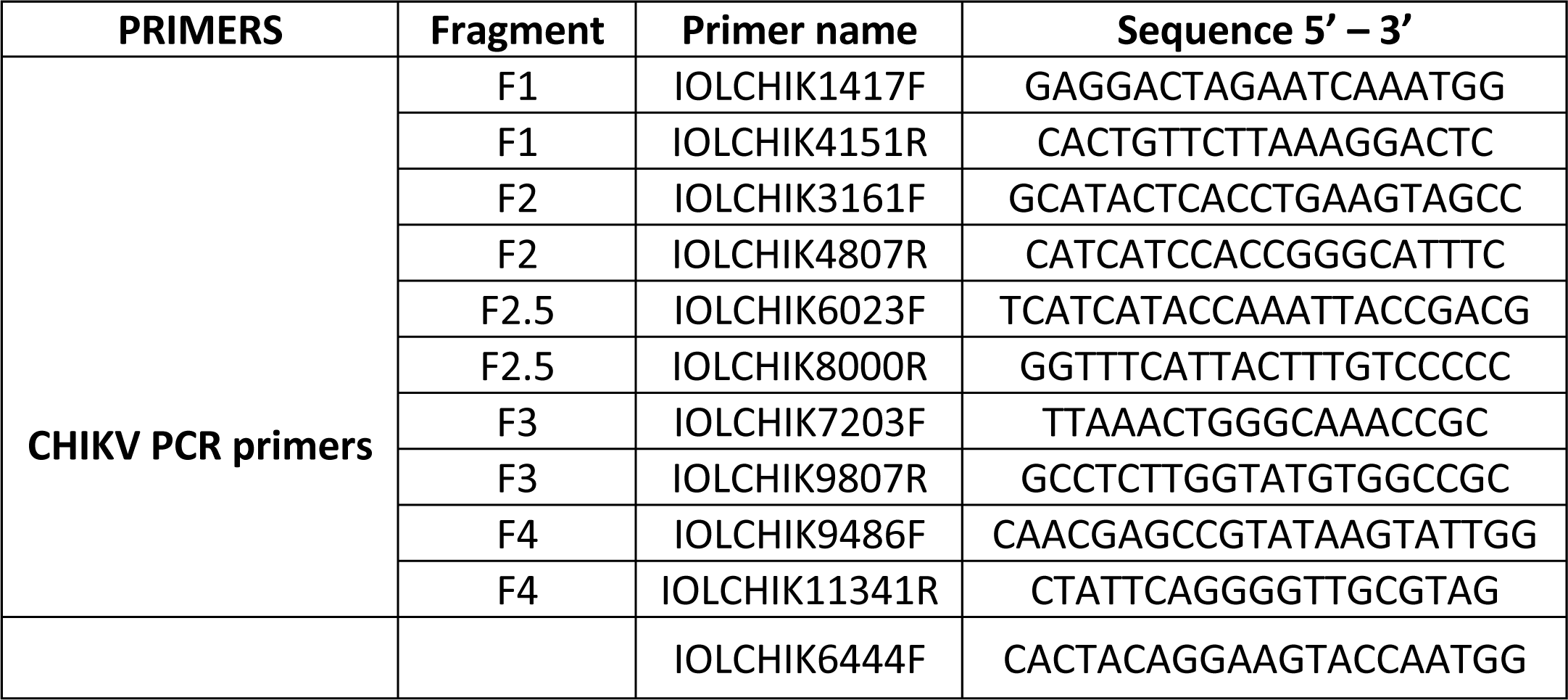
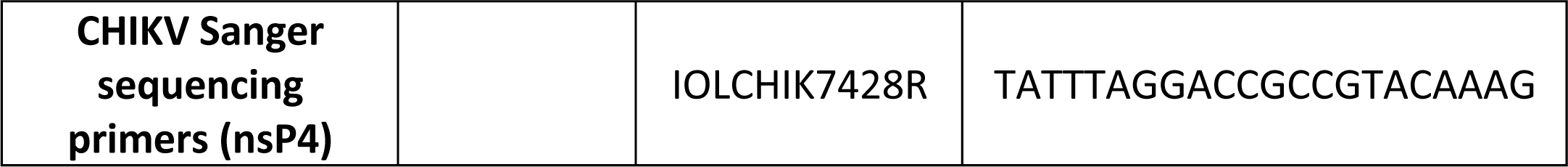
Primers used for Sanger sequencing.

### Deep sequencing library preparation

Deep sequencing library preparation and sequencing was performed at the NYU Genome Technology Center. The full length CHIKV genome was amplified as described above and fragments F1 / F2 / F2.5 / F3 and F4 of a given sample were mixed to obtain an equimolar amplicon mix spanning the whole genome. Each mix was then purified using the NucleoSpin Gel and PCR Clean-up kit by Macherey-Nagel (ref: 740609.50S). DNA was then eluted and quantified at a Nanodrop 2000 Spectrophotometer (ThermoScientific). A total of 25 ng of sample was put in each well of a 96-wp and adjusted with purified water to a final volume of 15 ΙL per well. Library preparations were made using the Nextera Flex Prep HT kit (Illumina) and samples were then sequenced on the Illumina NovaSeq X-Plus (300 cycles).

### Deep sequencing analysis

Sequencing raw fasta files were trimmed using Trimmomatic V0.39 and aligned using STAR version 2.7 (58) against the reference genome CHIKV strain La Reunion 06-049 (GenBank accession no. AM258994). The aligned SAM sequences were converted in BAM files, sorted, and indexed using Samtools which version v1.18 (59). Deduplication was then done by Picard.jar Mark Duplicates v3.1 (60). Variants were called using iVar (61). Minority variants at a given nucleotide position with a coverage depth of 500x or above (**Fig S5**) were selected with a minimum quality score of 20. An additional Fisher test was performed to check for strand bias and reads were excluded if the test turned true. Only variants present at least in technical or biological duplicates were retained. A conservative threshold of 0.01 (1%) was chosen. The RMSD was calculated on the common variants of each nsP4 variant according to Li. *et al*, 2010 ANDES: Statistical tools for the Analyses of Deep Sequencing (62) and represents the variance at each nucleotide position across the whole genome. All analysis code can be found on GitHub as the following web link: https://github.com/bbonaventure/CHIKV_fidelity/tree/main

### Protein structure analysis

The CHIKV replication complex structure containing the O’nyong’nyong virus nsP4 structure (91% similarity to CHIKV (21)) used in this paper is the one solved by Tan Y. B. *et al, Science Advances* (2022), PDB: 7Y38. The structure was visualized using PyMOL (version 2.5.2).

### Alphavirus sequence alignment and consensus sequence analysis

For nsP4 alignment and WebLogo generation, 12 representative members of the *Alphavirus* genus were selected based on Tan Y. B. *et al, Science Advances* (2022). Specifically, nsP4 protein sequences were first aligned using the multiple sequence alignment algorithm MUSCLE (EMBL-EMI). Following are their GenBank accession numbers: CHIKV: NC_004162, ONNV: AF079456.1, SFV: NC_003215, SINV: NC_001547, RRV: GQ433354.1, MAYV: NC_003417.1, BFV: MN689034.1, VEEV: L01442.2, EEEV: EF151502.1, WEEV: MN477208.1, EILV: NC_018615.1, GETV: NC_006558.1. Using the generated multiple sequence alignment of nsP4, a consensus sequence logo was designed via WebLogo with the default color scheme. The height of each stack of letters represents the sequence conservation at the indicated position and the height of each symbol within the stack represents its relative frequency.

### Statistical analysis

Statistical analysis was performed using GraphPad Prism (version 9.3.1). Mann-Whitney *t* test was performed against CHIKV wild-type condition, with *p* values < 0.05 considered significant. All experiments were done in two or three independent biological experiments and in two internal technical duplicates (details could be found in the legends of the figures). Error bars show +/- SEM or SD as specified in each figure’s legend.

## Data availability statement

Deep sequencing data have been deposited at the Sequence Read Archive (NIH) and is available under the following BioProject ID PRJNA1051608.

## Acknowledgments

We thank all the members of the Stapleford lab for their helpful comments and suggestions on this project. We thank Drs. Ludovic Desvignes, Dominick Papandrea and Alison Gilchrist for their support with the NYU ABSL3 high-containment laboratory and Dr. Yan Deng from the NYU Microscopy laboratory for his support with the use of the Zeiss LSM880 microscope. We also thank the Genomic Technology center (NYU, New York, RRID: SCR_017929) for the library preparation and for running the deep sequencing. This work was supported by a start-up package from the NYU Grossman School of Medicine, NIH/NIAID R01 AI162774-01 (K.A.S). MFM is supported by a Jan Vilcek/David Goldfarb Fellowship from the New York University Department of Microbiology, New York University (NYC, USA).

## Author’s contribution

Conceived and designed the experiments: MFM, OBP, KRG, KAS

Performed the experiments: MFM, NEM

Analyzed the data: MFM, NEM, BB, KAS

Contributed reagents/material/analysis tools: BB

Wrote the paper: MFM, KAS

## Supplementary information

**Supplemental Table 1: Minority variants found in this study.**

**S1 Fig. Alphaviruses nsP4 protein alignment.**

Twelve representative members of the *Alphavirus* genus (CHIKV, ONNV, SFV, SINV, RRV, MAYV, BFV, VEEV, EEEV, WEEV, EILV, GETV) were selected and nsP4 protein sequences were aligned using the multiple sequence alignment algorithm MUSCLE (EMBL-EMI). Each color represents amino acid with the same physicochemical properties (red: small hydrophobic including aromatic except Y; bleu: acidic; magenta: basic except H; green: hydroxyl, sulfhydryl, amine, glycine; grey: unusual amino acid). Symbols at the bottom of each amino acid indicates the conservation level of the position. An asterisk indicates positions which have a single, fully conserved residue. A colon indicates conservation between groups of strongly similar properties. A period indicates conservation between groups of weakly similar properties.

**S2 Fig. Intracellular viral proteins in mammalian cells.** Intracellular proteins from transfected BHK-21 cells from a first independent biological experiment (**A**) and a second independent biological experiment (**B**) were harvested at 48 hpt. Proteins were separated by SDS-PAGE, transferred to PVDF, and immunoblotted for the structural proteins capsid and E2 (top of a membrane), nsP4 (middle of a membrane) and the house-keeping gene actin (bottom of a membrane) with molecular weights on the left side as a reference. The nsP4 variants were split onto two distinct membranes, ran in the same tank, with all three controls, Mock, nsP4 WT and nsP4 GNN, were ran on each gel and the nsP4 variants organized from the N-ter (left of the membrane) to the C-ter (right of the membrane) of nsP4. The red star highlights the nsP4 variants that reverted in the specific experiment shown here.

**S3 Fig. Intracellular viral proteins in mosquito cells.** Intracellular proteins from transfected C6/36 from a first independent biological experiment (**A**) and a second biological experiment (**B**) were harvested at 48 hpt. Proteins were separated by SDS-PAGE, transferred to PVDF, and immunoblotted for the structural proteins Capsid and E2 (top of a membrane), nsP4 (middle of a membrane) and the house-keeping gene actin (bottom of a membrane) with molecular weights on the left side as a reference. The nsP4 variants were split onto two distinct membranes, ran in the same tank, with all three controls, Mock, nsP4 WT and nsP4 GNN, were ran on each gel and the nsP4 variants organized from the N-ter (left of the membrane) to the C- ter (right of the membrane) of nsP4. The red star highlights the nsP4 variants that reverted in the specific experiment shown here.

**S4 Fig. Extracellular viral proteins in mosquito cells.** Supernatants from C6/36 cells transfected with *in vitro* transcribed full-length CHIKV nsP4 variants that did not show any infectious viral particles, but intracellular viral proteins were harvested at 48 hpt. Extracellular proteins were purified with Amicon columns. Denatured proteins were subjected to a SDS-PAGE and western blotting to for the presence of the structural proteins capsid (top), E2 (middle) and E1 (bottom). A first independent biological experiment is shown in (**A**) and a second one in (**B**). The corresponding Coomassie blue is presented at the bottom of each Western Blot.

**S5 Fig. Immunofluorescence negative controls and additional variant images. (A)** Mock transfected BHK-21 cells and **(B)** BHK-21 cells transfected with the replication-dead nsP4 GNN variant stained for nsP4 and DAPI. **(C and D)** Mock transfected C6/36 cells or C6/36 cells transfected with nsP4 variants and stained for nsP4, capsid, E2, and actin.

**S6 Fig. Deep sequencing depth in mammalian and mosquito cells.** BHK-21 cells (**A, B, C and D**) and C6/36 (**E, F, G and H**) deep sequencing depth (total reads) is depicted per nucleotide position along the genome. Shown in BHK-21 cells is nsP4 WT (**A**), nsp4 C483Y (**B**), nsP4 W486F (**C**), nsP4 W486Y (**D**) and in C6/36 cells is nsP4 WT (**E**), nsP4 L383F (**F**), nsP4 C483Y (**G**) and nsP4 W486Y (**H**). Each color represents one independent biological experiment or replicate.

**S7 Fig. Deep sequencing analysis of nsP4 W486F in mammalian cells.** Supernatants from transfection duplicates of the genetically stable W486F variant in BHK-21 cells (corresponding to the Fig 2) were subjected to deep sequencing. The bioinformatic analysis was performed using a conservative threshold of 1% for the variant calling. (**A**) Variant frequency by nucleotide position along the genome for the nsP4 W486F variant. Nonsynonymous mutations are colored in grey, synonymous mutations in salmon and non-coding mutations in blue. The variants of interest nsP2 I502V and nsP4 V209I are highlighted in a dashed circle, respectively colored in green or red. A schematic of CHIKV genome is depicted at the bottom of the graph. (**B**) Absolute number of minority variants showing nsP4 WT, nsP4 C483Y, nsP4 W486F and nsP4 W486Y variants. The corresponding dN/dS ratio is depicted in (**C**) and the Root Mean Square Deviation (RMSD) in (**D**). The RMSD is a measure of the average diversity generated by the nsP4 variants at each position across the genome. For all panels, a Mann-Whitney *t* test was performed against nsP4 WT but no data reached statistical significance. Graphs show the average and SEM of the two technical replicates. The percentage of transitions and transversions events is shown in (**E**). Analysis of the specific nucleotide changes frequency corresponding to nsP4 WT, nsP4 C483Y, nsP4 W486F and nsP4 W486Y in BHK-21 cells is presented in (**F**). Each substitution type is associated to a given color: red for U to C, orange for G to A, yellow for G to U, grey for C to A, light blue for A to G, pink for A to U, dark blue for C to U, purple for A to C, mauve for U to A, medium blue for G to C, green for C to G and brown for U to G. (**G**) Zoom on the core replicase 3D structure (PDB: 7Y38) and the potential interactions between nsP4 V209 (depicted in pink) and residues from a nsP2 loop (depicted in turquoise). W486 is depicted in brown. Potential interaction between the nsP4 V209 side chain and either nsP2 K396 or nsP2 E399 is presented. For all panels, a Mann-Whitney *t* test was performed against nsP4 WT but no data reached statistical significance. Graphs show the average and SEM of three biological independent experiments.

## Notes

### Competing Interest Statement

The authors have declared no competing interest.

## References

1. Hahn MB, Eisen L, McAllister J, Savage HM, Mutebi JP, Eisen RJ. Updated Reported Distribution of Aedes (Stegomyia) aegypti and Aedes (Stegomyia) albopictus (Diptera: Culicidae) in the United States, 1995–2016. Journal of Medical Entomology. 2017 Sep 1;54(5):1420–4.

2. Schwartz O, Albert ML. Biology and pathogenesis of chikungunya virus. Nat Rev Microbiol. 2010 Jul;8(7):491–500.

3. Feldstein LR, Ellis EM, Rowhani-Rahbar A, Hennessey MJ, Staples JE, Halloran ME, et al. Estimating the cost of illness and burden of disease associated with the 2014–2015 chikungunya outbreak in the U.S. Virgin Islands. Maheu-Giroux M, editor. PLoS Negl Trop Dis. 2019 Jul 19;13(7):e0007563.

4. Costa LB, Barreto FKDA, Barreto MCA, Santos THPD, Andrade MDMOD, Farias LABG, et al. Epidemiology and Economic Burden of Chikungunya: A Systematic Literature Review. TropicalMed. 2023 May 31;8(6):301.

5. Kril V, Aïqui-Reboul-Paviet O, Briant L, Amara A. New Insights into Chikungunya Virus Infection and Pathogenesis. Annu Rev Virol. 2021 Sep 29;8(1):327–47.

6. Ahola T, Kääriäinen L. Reaction in alphavirus mRNA capping: formation of a covalent complex of nonstructural protein nsP1 with 7-methyl-GMP. Proceedings of the National Academy of Sciences. 1995 Jan 17;92(2):507–11.

7. Gottipati K, Woodson M, Choi KH. Membrane binding and rearrangement by chikungunya virus capping enzyme nsP1. Virology. 2020 May;544:31–41.

8. Jones R, Bragagnolo G, Arranz R, Reguera J. Capping pores of alphavirus nsP1 gate membranous viral replication factories. Nature. 2021 Jan 28;589(7843):615–9.

9. Tan YB, Lello LS, Liu X, Law YS, Kang C, Lescar J, et al. Crystal structures of alphavirus nonstructural protein 4 (nsP4) reveal an intrinsically dynamic RNA-dependent RNA polymerase fold. Nucleic Acids Research. 2022 Jan 25;50(2):1000–16.

10. Vasiljeva L, Merits A, Auvinen P, Kääriäinen L. Identification of a Novel Function of the AlphavirusCapping Apparatus. Journal of Biological Chemistry. 2000 Jun;275(23):17281–7.

11. Russo AT, White MA, Watowich SJ. The Crystal Structure of the Venezuelan Equine Encephalitis Alphavirus nsP2 Protease. Structure. 2006 Sep;14(9):1449–58.

12. Das PK, Merits A, Lulla A. Functional Cross-talk between Distant Domains of Chikungunya Virus Non-structural Protein 2 Is Decisive for Its RNA-modulating Activity*. Journal of Biological Chemistry. 2014 Feb 28;289(9):5635–53.

13. Stapleford KA, Rozen-Gagnon K, Das PK, Saul S, Poirier EZ, Blanc H, et al. Viral Polymerase-Helicase Complexes Regulate Replication Fidelity To Overcome Intracellular Nucleotide Depletion. Journal of Virology. 2015 Oct 22;89(22):11233–44.

14. Abraham R, Hauer D, McPherson RL, Utt A, Kirby IT, Cohen MS, et al. ADP-ribosyl– binding and hydrolase activities of the alphavirus nsP3 macrodomain are critical for initiation of virus replication. Proc Natl Acad Sci USA [Internet]. 2018 Oct 30 [cited 2023 Oct 24];115(44). Available from: https://pnas.org/doi/full/10.1073/pnas.1812130115

15. Meshram CD, Agback P, Shiliaev N, Urakova N, Mobley JA, Agback T, et al. Multiple Host Factors Interact with the Hypervariable Domain of Chikungunya Virus nsP3 and Determine Viral Replication in Cell-Specific Mode. López S, editor. J Virol. 2018 Aug 15;92(16):e00838–18.

16. Tomar S, Hardy RW, Smith JL, Kuhn RJ. Catalytic Core of Alphavirus Nonstructural Protein nsP4 Possesses Terminal Adenylyltransferase Activity. Journal of Virology. 2006 Oct 15;80(20):9962–9.

17. Coffey LL, Beeharry Y, Bordería AV, Blanc H, Vignuzzi M. Arbovirus high fidelity variant loses fitness in mosquitoes and mice. Proceedings of the National Academy of Sciences. 2011 Sep 20;108(38):16038–43.

18. Rozen-Gagnon K, Stapleford KA, Mongelli V, Blanc H, Failloux AB, Saleh MC, et al. Alphavirus Mutator Variants Present Host-Specific Defects and Attenuation in Mammalian and Insect Models. Kramer LD, editor. PLoS Pathog. 2014 Jan 16;10(1):e1003877.

19. Tan YB, Chmielewski D, Law MCY, Zhang K, He Y, Chen M, et al. Molecular architecture of the Chikungunya virus replication complex. Science Advances. 2022 Nov 30;8(48):eadd2536.

20. Freire MCLC, Basso LGM, Mendes LFS, Mesquita NCMR, Mottin M, Fernandes RS, et al. Characterization of the RNA-dependent RNA polymerase from Chikungunya virus and discovery of a novel ligand as a potential drug candidate. Sci Rep. 2022 Jun 22;12(1):10601.

21. Solignat M, Gay B, Higgs S, Briant L, Devaux C. Replication cycle of chikungunya: A re-merging arbovirus. Virology. 2009 Oct 25;393(2):183–97.

22. Gnädig NF, Beaucourt S, Campagnola G, Bordería AV, Sanz-Ramos M, Gong P, et al. Coxsackievirus B3 mutator strains are attenuated in vivo. Proceedings of the National Academy of Sciences. 2012 Aug 21;109(34):E2294–303.

23. Campagnola G, McDonald S, Beaucourt S, Vignuzzi M, Peersen OB. Structure-Function Relationships Underlying the Replication Fidelity of Viral RNA-Dependent RNA Polymerases. Kirkegaard K, editor. J Virol. 2015 Jan;89(1):275–86.

24. Martinez MG, Kielian M. Intercellular Extensions Are Induced by the Alphavirus Structural Proteins and Mediate Virus Transmission. Whelan SPJ, editor. PLoS Pathog. 2016 Dec 15;12(12):e1006061.

25. Yin P, Davenport BJ, Wan JJ, Kim AS, Diamond MS, Ware BC, et al. Chikungunya virus cell-to-cell transmission is mediated by intercellular extensions in vitro and in vivo. Nat Microbiol. 2023 Sep;8(9):1653–67.

26. Weiss CM, Liu H, Riemersma KK, Ball EE, Coffey LL. Engineering a fidelity-variant live-attenuated vaccine for chikungunya virus. npj Vaccines. 2020 Oct 14;5(1):1–13.

27. Lyons DM, Lauring AS. Evidence for the Selective Basis of Transition-to-Transversion Substitution Bias in Two RNA Viruses. Molecular Biology and Evolution. 2017 Dec 1;34(12):3205–15.

28. Gong P, Peersen OB. Structural basis for active site closure by the poliovirus RNA- dependent RNA polymerase. Proceedings of the National Academy of Sciences. 2010 Dec 28;107(52):22505–10.

29. Coffey LL, Forrester N, Tsetsarkin K, Vasilakis N, Weaver SC. Factors shaping the adaptive landscape for arboviruses: implications for the emergence of disease. Future Microbiology. 2013 Feb;8(2):155–76.

30. Kayikci M, Venkatakrishnan AJ, Scott-Brown J, Ravarani CNJ, Flock T, Babu MM. Visualization and analysis of non-covalent contacts using the Protein Contacts Atlas. Nat Struct Mol Biol. 2018 Feb;25(2):185–94.

31. Liu B, Dai R, Tian CJ, Dawson L, Gorelick R, Yu XF. Interaction of the Human Immunodeficiency Virus Type 1 Nucleocapsid with Actin. J Virol. 1999 Apr;73(4):2901–8.

32. Thomas A, Mariani-Floderer C, López-Huertas MR, Gros N, Hamard-Péron E, Favard C, et al. Involvement of the Rac1-IRSp53-Wave2-Arp2/3 Signaling Pathway in HIV-1 Gag Particle Release in CD4 T Cells. Journal of Virology. 2015 Jul 21;89(16):8162–81.

33. Bedi S, Ono A. Friend or Foe: The Role of the Cytoskeleton in Influenza A Virus Assembly. Viruses. 2019 Jan;11(1):46.

34. Zhang Y, Gao W, Li J, Wu W, Jiu Y. The Role of Host Cytoskeleton in Flavivirus Infection. Virol Sin. 2019 Feb;34(1):30–41.

35. Dibsy R, Bremaud E, Mak J, Favard C, Muriaux D. HIV-1 diverts cortical actin for particle assembly and release. Nat Commun. 2023 Oct 31;14(1):6945.

36. Kathuria SV, Chan YH, Nobrega RP, Özen A, Matthews CR. Clusters of isoleucine, leucine, and valine side chains define cores of stability in high-energy states of globular proteins: Sequence determinants of structure and stability. Protein Science. 2016;25(3):662–75.

37. Shu B, Gong P. Structural basis of viral RNA-dependent RNA polymerase catalysis and translocation. Proceedings of the National Academy of Sciences. 2016 Jul 12;113(28):E4005–14.

38. Vignuzzi M, López CB. Defective viral genomes are key drivers of the virus–host interaction. Nat Microbiol. 2019 Jul;4(7):1075–87.

39. Poirier EZ, Mounce BC, Rozen-Gagnon K, Hooikaas PJ, Stapleford KA, Moratorio G, et al. Low-Fidelity Polymerases of Alphaviruses Recombine at Higher Rates To Overproduce Defective Interfering Particles. Journal of Virology. 2016 Feb 11;90(5):2446–54.

40. Mendes A, Kuhn RJ. Alphavirus Nucleocapsid Packaging and Assembly. Viruses. 2018 Mar;10(3):138.

41. Jose J, Taylor AB, Kuhn RJ. Spatial and Temporal Analysis of Alphavirus Replication and Assembly in Mammalian and Mosquito Cells. mBio. 2017 Feb 14;8(1):10.1128/mbio.02294-16.

42. Lulla V, Kim DY, Frolova EI, Frolov I. The Amino-Terminal Domain of Alphavirus Capsid Protein Is Dispensable for Viral Particle Assembly but Regulates RNA Encapsidation through Cooperative Functions of Its Subdomains. Journal of Virology. 2013 Nov 15;87(22):12003–19.

43. White CL, Thomson M, Dimmock NJ. Deletion Analysis of a Defective Interfering Semliki Forest Virus RNA Genome Defines a Region in the nsP2 Sequence That Is Required for Efficient Packaging of the Genome into Virus Particles. J Virol. 1998 May;72(5):4320–6.

44. Kim DY, Atasheva S, Frolova EI, Frolov I. Venezuelan Equine Encephalitis Virus nsP2 Protein Regulates Packaging of the Viral Genome into Infectious Virions. J Virol. 2013 Apr 15;87(8):4202–13.

45. Rana J, Rajasekharan S, Gulati S, Dudha N, Gupta A, Chaudhary VK, et al. Network mapping among the functional domains of Chikungunya virus nonstructural proteins. Proteins: Structure, Function, and Bioinformatics. 2014;82(10):2403–11.

46. Forsell K, Suomalainen M, Garoff H. Structure-function relation of the NH2-terminal domain of the Semliki Forest virus capsid protein. Journal of Virology. 1995 Mar;69(3):1556– 63.

47. Rupp JC, Jundt N, Hardy RW. Requirement for the Amino-Terminal Domain of Sindbis Virus nsP4 during Virus Infection. J Virol. 2011 Apr;85(7):3449–60.

48. Denison MR, Graham RL, Donaldson EF, Eckerle LD, Baric RS. Coronaviruses: An RNA proofreading machine regulates replication fidelity and diversity. RNA Biology. 2011 Mar;8(2):270–9.

49. Smith EC, Blanc H, Vignuzzi M, Denison MR. Coronaviruses Lacking Exoribonuclease Activity Are Susceptible to Lethal Mutagenesis: Evidence for Proofreading and Potential Therapeutics. Diamond MS, editor. PLoS Pathog. 2013 Aug 15;9(8):e1003565.

50. Andzhaparidze OG, Bogomolova NN, Boriskin YS, Bektemirova MS, Drynov ID. Comparative study of rabies virus persistence in human and hamster cell lines. Journal of Virology. 1981 Jan;37(1):1–6.

51. Brackney DE, Scott JC, Sagawa F, Woodward JE, Miller NA, Schilkey FD, et al. C6/36 Aedes albopictus Cells Have a Dysfunctional Antiviral RNA Interference Response. O’Neill SL, editor. PLoS Negl Trop Dis. 2010 Oct 26;4(10):e856.

52. Scott JC, Brackney DE, Campbell CL, Bondu-Hawkins V, Hjelle B, Ebel GD, et al. Comparison of Dengue Virus Type 2-Specific Small RNAs from RNA Interference-Competent and –Incompetent Mosquito Cells. O’Neill SL, editor. PLoS Negl Trop Dis. 2010 Oct 26;4(10):e848.

53. Roy A, Kucukural A, Zhang Y. I-TASSER: a unified platform for automated protein structure and function prediction. Nat Protoc. 2010 Apr;5(4):725–38.

54. Schwede T. SWISS-MODEL: an automated protein homology-modeling server. Nucleic Acids Research. 2003 Jul 1;31(13):3381–5.

55. Kelley LA, Mezulis S, Yates CM, Wass MN, Sternberg MJE. The Phyre2 web portal for protein modeling, prediction and analysis. Nat Protoc. 2015 Jun;10(6):845–58.

56. Peersen O. A Comprehensive Superposition of Viral Polymerase Structures. Viruses. 2019 Aug 13;11(8):745.

57. Carrau L, Rezelj VV, Noval MG, Levi LI, Megrian D, Blanc H, et al. Chikungunya Virus Vaccine Candidates with Decreased Mutational Robustness Are Attenuated *In Vivo* and Have Compromised Transmissibility. Dermody TS, editor. J Virol. 2019 Sep 15;93(18):e00775-19.

58. Dobin A, Davis CA, Schlesinger F, Drenkow J, Zaleski C, Jha S, et al. STAR: ultrafast universal RNA-seq aligner. Bioinformatics. 2013 Jan 1;29(1):15–21.

59. Li H, Handsaker B, Wysoker A, Fennell T, Ruan J, Homer N, et al. The Sequence Alignment/Map format and SAMtools. Bioinformatics. 2009 Aug 15;25(16):2078–9.

60. Picard Tools - By Broad Institute [Internet]. [cited 2023 Nov 20]. Available from: https://broadinstitute.github.io/picard/

61. Grubaugh ND, Gangavarapu K, Quick J, Matteson NL, De Jesus JG, Main BJ, et al. An amplicon-based sequencing framework for accurately measuring intrahost virus diversity using PrimalSeq and iVar. Genome Biology. 2019 Jan 8;20(1):8.

62. Li K, Venter E, Yooseph S, Stockwell TB, Eckerle LD, Denison MR, et al. ANDES: Statistical tools for the ANalyses of DEep Sequencing. BMC Research Notes. 2010 Jul 15;3(1):199.

